# Scedar: a scalable Python package for single-cell RNA-seq exploratory data analysis

**DOI:** 10.1101/375196

**Authors:** Yuanchao Zhang, Man S. Kim, Erin R. Reichenberger, Ben Stear, Deanne M. Taylor

## Abstract

In single-cell RNA-seq (scRNA-seq) experiments, the number of individual cells has increased exponentially, and the sequencing depth of each cell has decreased significantly. As a result, analyzing scRNA-seq data requires extensive considerations of program efficiency and method selection. In order to reduce the complexity of scRNA-seq data analysis, we present scedar, a scalable Python package for scRNA-seq exploratory data analysis. The package provides a convenient and reliable interface for performing visualization, imputation of gene dropouts, detection of rare transcriptomic profiles, and clustering on large-scale scRNA-seq datasets. The analytical methods are efficient, and they also do not assume that the data follow certain statistical distributions. The package is extensible and modular, which would facilitate the further development of functionalities for future requirements with the open-source development community. The scedar package is distributed under the terms of the MIT license at https://pypi.org/project/scedar.

## 1 Introduction

Cost-effective large-scale transcriptomic profiling of individual cells is enabled by the development of microfluidic, nanodroplet, and massively parallel sequencing technologies. Using these technologies, single-cell RNA-seq (scRNA-seq) experiments usually generate transcriptomic profiles of thousands to millions of individual cells (Svensson, Vento-Tormo, and Teichmann 2018). Therefore, scRNA-seq has become more commonly used to either study specific biological questions or comprehensively profile certain tissues or organisms (Filbin et al. 2018; Cao et al. 2017; Regev et al. 2017).

Analyses of scRNA-seq datasets therefore require efficient computational programs and sophisticated statistical methods. The programs should be able to manage memory efficiently, exploit multiple cores of the processing units, and handle errors and exceptions gracefully. The statistical methods must be able to function against high dimensionality, low signal-to-noise ratio, and different characteristics of data generated from different technologies and protocols (Kiselev, Andrews, and Hemberg 2019; Dueck et al. 2016; Ziegenhain et al. 2017). Such requirements can become a barrier between experimental design and biological interpretations of the results.

In order to be scalable, methods have been designed to minimize the usage of hardware resources, so that a large-scale scRNA-seq dataset can be analyzed using a desktop computer, such as Seurat v3.0 (Butler et al. 2018) and Scanpy (Wolf, Angerer, and Theis 2018). Seurat is an R package providing visualization and robust statistical methods to explore and interpret the heterogeneity of the dataset. Scanpy is a Python package providing efficient reimplementations of pre-existing statistical methods and analytical workflows, which can be used to perform general exploratory data analysis and special inferences.

Here, we attempt to achieve the scalability of scRNA-seq data analysis through an alternative approach, which is to enable the user to exploit powerful analytical methods using modern high-performance computing architectures (Zaharia et al. 2016; Dean and Ghemawat 2008; Abadi et al. 2016), such as servers and clusters with large amount of memory, multiple central and graphical processing unit cores, and solid-state drives (SSD) with higher read and write speed than traditional hard disk drive (HDD). Such computational resources have been made easily accessible by cloud computing services like Amazon Web Services and Microsoft Azure. Applying this approach, we designed the analytical methods to be able to run in parallel processes with minimized memory overhead.

Additionally, we also aim to develop robust exploratory data analysis (EDA) methods, so that they could be applied to various datasets generated by different experimental designs, technologies, and protocols. Single-cell RNA-seq datasets have distinct statistical characteristics if generated from different scRNA-seq technologies and platforms (Zappia, Phipson, and Oshlack 2017; Dueck et al. 2016; Ziegenhain et al. 2017), such as SMARTer (Pollen et al. 2014), Drop-seq (Macosko et al. 2015) and 10x Genomics GemCode (Zheng et al. 2017). In order to avoid assumptions on the statistical distributions of the datasets, we incorporated efficient implementations of machine learning methods into the data exploration process. Through extensive exploration, the statistical properties of the data could be observed and used to guide the selection of appropriate methods for biological interpretation (Zappia, Phipson, and Oshlack 2018).

Therefore, we developed a scalable and reliable Python package, **<u>s</u>**ingle-**<u>c</u>**ell **<u>e</u>**xploratory **<u>d</u>**ata analysis for **<u>R</u>**NA-seq (scedar), to facilitate the exploration of large-scale scRNA-seq datasets. Scedar provides analytical routines for visualization, gene dropout imputation, rare transcriptomic profile detection, clustering, and identification of cluster separating genes. The visualization methods are integrated with the efficient scRNA-seq data structures to provide intuitive, convenient, and flexible plotting interfaces. We implemented methods to impute gene dropouts (Supplementary Algorithm S1) and detect rare transcriptomic profiles (Supplementary Algorithm S2) based on the k-nearest neighbor (KNN) algorithm. The detected rare transcriptomic profiles could be compared with their nearest neighbors in detail to identify rare cell states or types. For clustering analysis, we provide a novel cell clustering algorithm named MIRAC (Algorithm 1), minimum description length (**<u>M</u>**DL) **<u>i</u>**teratively **<u>r</u>**egularized **<u>a</u>**gglomerative **<u>c</u>**lustering. In order to identify genes that are able to distinguish the clusters, we provide a method using a sparsity-aware gradient boosted tree system, XGBoost (Chen and Guestrin 2016).

## 2 Methods

### 2.1 scedar package design

We designed scedar in an object-oriented manner for quickly exploring large-scale scRNA-seq transcription level matrices on a remote server utilizing parallel computing techniques, in order to provide a robust and extensible platform for EDA, rather than surpassing the analytical performance of any of ≥ 275 existing scRNA-seq data analysis methods (Zappia, Phipson, and Oshlack 2018). The application programming interface (API) is designed to be intuitive to users familiar with the R programming language. The standard analysis workflow is to explore the dataset, cluster cells, and identify cluster separating genes (**Figure 1**), which is implemented as four main modules: data structure, KNN, clustering and visualization.

**Figure 1.**
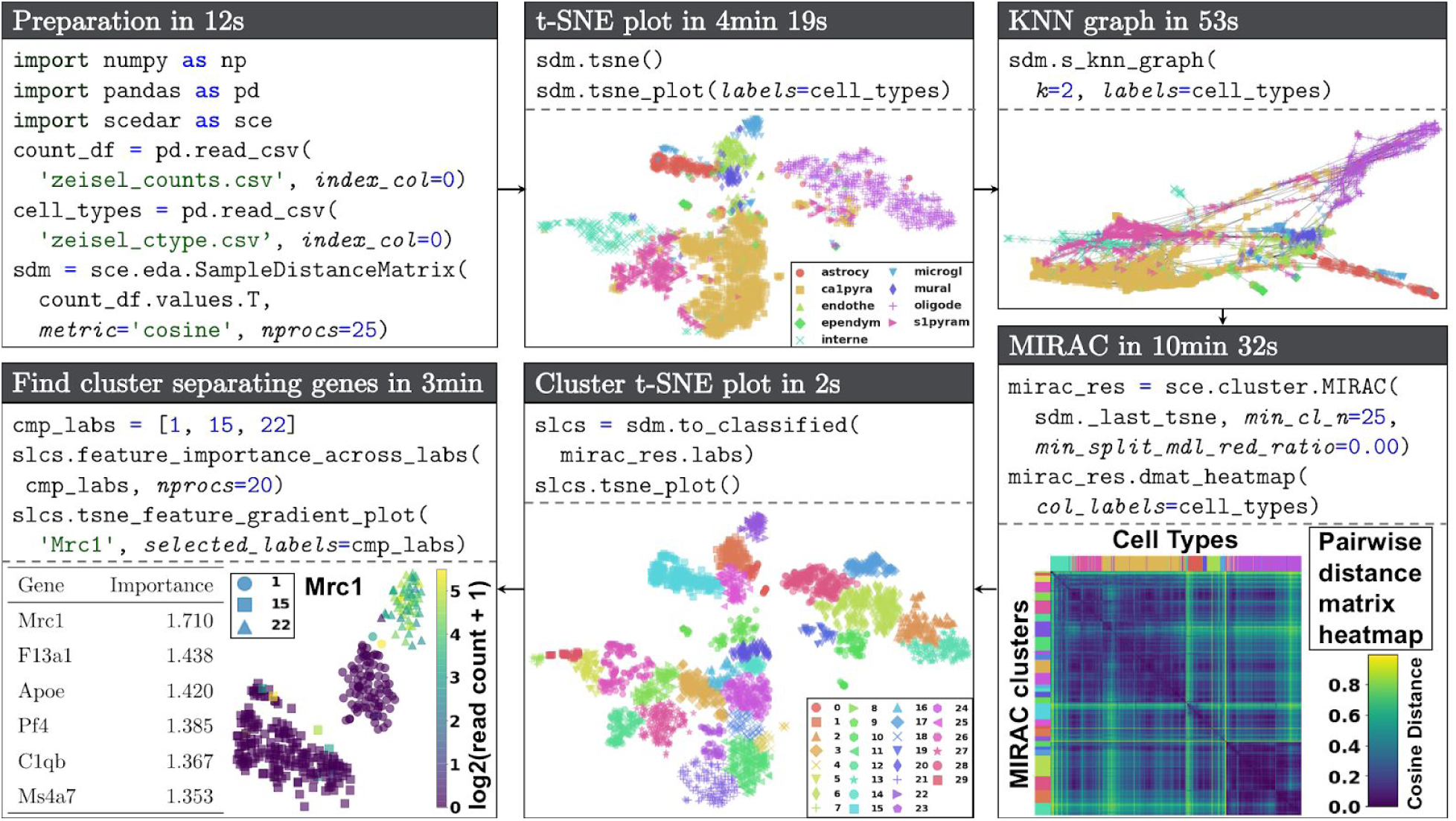
Demo of scedar. Workflow of using scedar to analyze an scRNA-seq dataset with 3005 mouse brain cells and 19,972 genes generated using the STRT-Seq UMI protocol by Zeisel *et al.* (Zeisel et al. 2015). Procedures and parameters that are not directly related to data analysis are omitted. The full version of the demo is available at https://github.com/logstar/scedar/tree/master/docs/notebooks.

The core data structure stores each transcription level matrix as a standalone sparse matrix or full array instance, and it can easily be extended to support customized analytical procedures. The built-in extensions include common procedures like pairwise distance computation, Principal Component Analysis (PCA), t-SNE (Maaten and Hinton 2008), UMAP (McInnes and Healy 2018), and k-nearest neighbor graph (Jacomy et al. 2014). We optimized time and memory efficiency with the following design patterns: parallel computing, lazy loading, caching and copy-on-write.

The KNN and clustering modules utilize the data structure and parallel computing to efficiently perform analytical procedures, and the results are stored in the data structure for further reference.

The visualization module contains plotting methods optimized for large datasets, especially for plotting heatmaps. For example, it takes less than two minutes to generate a heatmap image from a 50,000 × 20,000 matrix containing random standard normal entries.

Preprocessing is implemented as selection and transformation routines of the core data structure, which is not a focus of scedar. The package is designed to identify the necessity of certain preprocessing procedures by extensively exploring the original state of the data. Although preprocessing, such as batch effect correction and normalization, could facilitate the detection of biological variances between cells, applying preprocessing methods correctly requires careful validation of their assumptions, otherwise they may introduce unwanted bias or variability (Hicks and Irizarry 2015).

### 2.2 Minimum description length iteratively regulated agglomerative clustering

Minimum description length (MDL) iteratively regulated agglomerative clustering (MIRAC) extends hierarchical agglomerative clustering (HAC) (Müllner 2011) in a divide and conquer manner for scRNA-seq data. Input with raw or dimensionality reduced scRNA-seq data, MIRAC starts with one round of bottom-up HAC to build a tree with optimal linear leaf ordering (Bar-Joseph, Gifford, and Jaakkola 2001), and the tree is then divided into small sub-clusters, which are further merged iteratively into clusters. Because each individual cluster becomes more homogenous with higher number of clusters, the iterative merging process is regularized with the MDL principle (Hansen and Yu 2001). The asymptotic time complexity of the MIRAC algorithm is O(n^4^ + mn^2^), where n is the number of samples, and m is the number of features. The space complexity is O(n^2^ + mn). Relevant mathematical theories and notations of MDL are briefly described in the following section. The pseudo-code of MIRAC is shown in **Algorithm 1**.

#### Algorithm 1: Minimum description length iteratively regulated agglomerative clustering

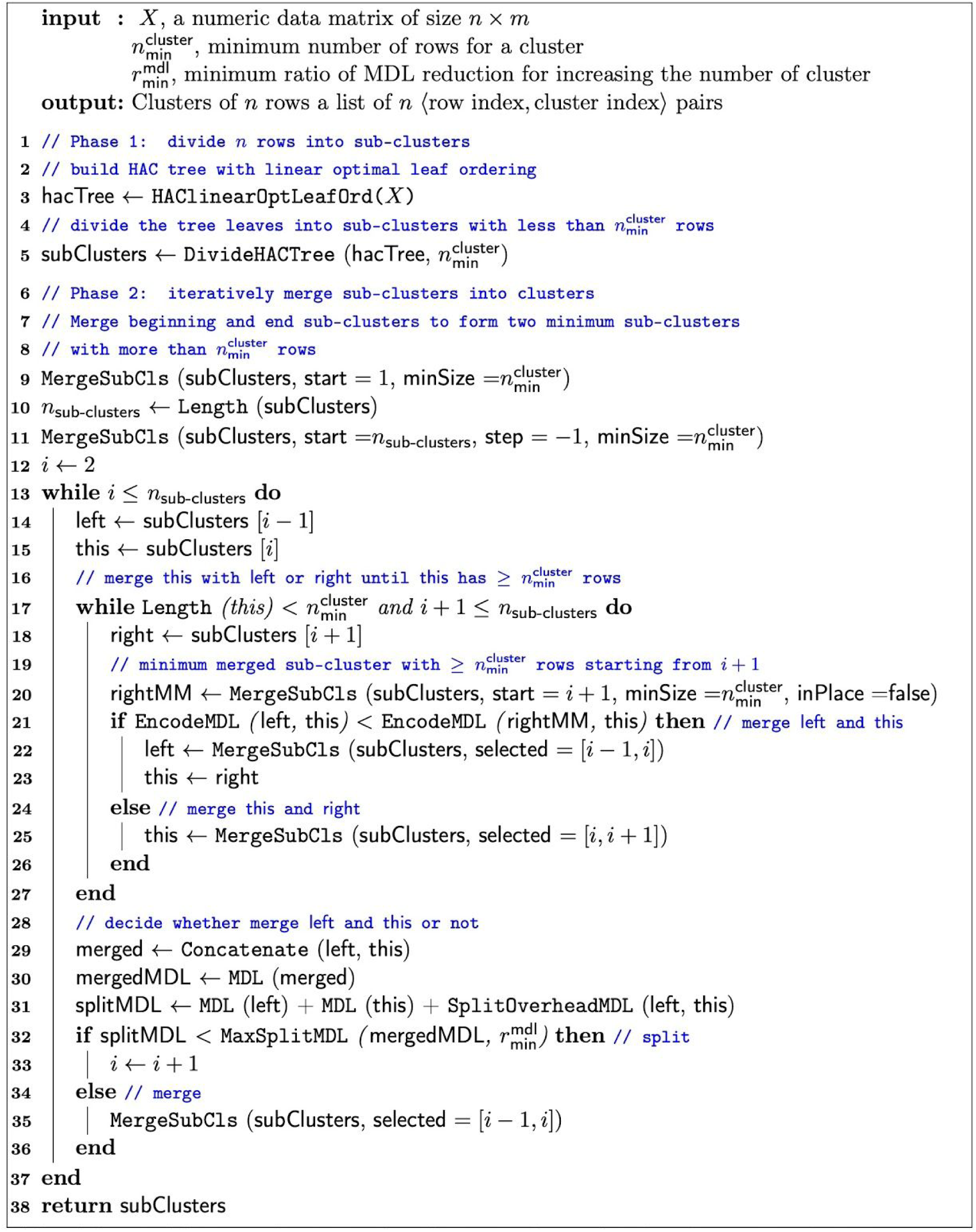

Comparing to HAC, MIRAC is not designed to be faster but rather to improve the cluster robustness. The asymptotic time complexity of MIRAC is the same as HAC with optimal leaf ordering. However, the use of MDL rather than deterministic similarity metrics, improves the noise tolerance by estimating similarity with probabilistic models to give more weight on signal and less weight on noise, assuming that the signal is stronger than noise.

#### 2.2.1 Rationale

The rationale behind MIRAC is to reduce the number of cell partitions that need to be evaluated and find a good one among them.

The number of all possible partitions of a set with n elements is the Bell number, which could be computed with the following recurrence 

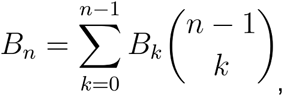

where *n* ≥ 1 and *B*_0_ = 1 (Wilf 2005).It is computationally intractable to compute the code lengths of *B*_*n*_ partitions, so we reduced the number of cell partitions to evaluate with the following steps:

1. We only evaluate the partitions of *n* cells that consecutively divide the optimal HAC tree *T* leaf ordering. The HAC tree with optimal leaf ordering maximizes the sum of similarity measurements between adjacent items in the leaf ordering (Bar-Joseph, Gifford, and Jaakkola 2001), which could be computed with an algorithm developed by Bar-Joseph *et al*. that runs in time O(n^4^). The number of partitions that consecutively divide an ordering is the same as the number of compositions of *n*, which is *C*_*n*_ = 2^*n*−1^. Because this number still grows exponentially with regard to *n*, it is necessary to further reduce the set of partitions for evaluation.
2. Within step 1 partitions, we only evaluate those with all clusters having ≥ *t* cells. The number of such partitions is the same as the compositions of *n* with all summands ≥ *t*, which is

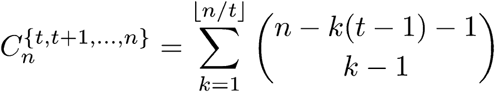

given by (Abramson 1976). The growth rate of 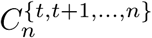 with regard to *n* is smaller, but we still need to reduce it further. For example, 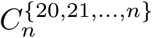 for *n* in an ⟨20, 40, 60, 80, 100, 120, 140, 160, 180, 200⟩ are ⟨1, 2, 23, 274, 2695, 24941, 232016, 2184520, 20628613, 194570810⟩, and the values of 2^*n*^ are ⟨1048576, 1099511627776, …, 1.607 × 10^60^⟩.
3. Within step 2 partitions, we only evaluate the ones that could be generated by merging adjacent clusters of a specific partition 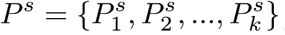, where *P*^*s*^ is generated by recursively divide the root HAC tree *T* until the partitioned subtree has ≤ *t* − 1 leaves. Thus, ⌈*n*/(*t* − 1) ⌉ ≤ *k* ≤ *n*. Let *M*(*P*^*s*^) denote the number of step 2 partitions that could be generated by merging adjacent clusters of *P*^*s*^. The upper bound of *M*(*P*^*s*^) is the same as the number of step 2 partitions, which could be reached when *k* = *n*. The lower bound of *M*(*P*^*s*^) is not straightforward, since the merged partitions should have all clusters with ≥ *t* cells.

In order to find a good cell partition *P*^*g*^ in the subset of all possible ones, we iteratively inspect each cluster of *P*^*s*^ and merge it with either its left or right adjacent cluster according to the similarity determined by the two-stage MDL scheme described in the following sections. The procedure has the following steps:

1. Merge *P*^*s*^ clusters in the beginning of the optimal ordering until the merged set contains ≥ *t* cells (**line 9 Algorithm 1**).
2. Similarly, merge *P*^*s*^ clusters at the end of the optimal ordering until the merged set contains ≥ *t* cells (**line 11 Algorithm 1**).
3. For every cluster 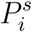 in the middle, determine the similarity between 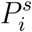 and the cluster on the left and right, and merge 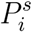 with the more similar cluster (**line 20 - 26 Algorithm 1**). Although the cluster on the left 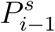 always has 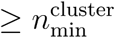 cells, the cluster on the right 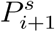 may have a minimum of one cell, so that the determined similarity between 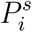 and 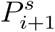 is sensitive to noise. In order to improve the robustness of similarity comparison, instead of determining the similarity between 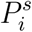 and 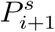, we determine the similarity between 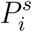 and 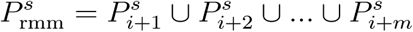 (**line 20 Algorithm 1**), where 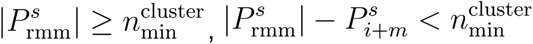, and rmm is the shorthand for right minimax.
4. Once 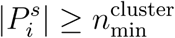, determine the similarity between 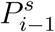 and 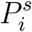, and merge them if their similarity is above a certain threshold, otherwise split them (**line 32 - 36 Algorithm 1**). The code length of merged 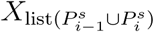 is

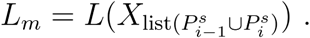 The code length of divided 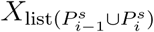 is

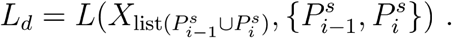 If 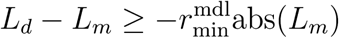, merge 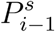 and 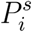, otherwise split them. This conditional statement generalizes the situations where *L*_*m*_ is either non-negative or negative.
5. Continue inspecting the next cluster in *P*^*s*^.
6. After inspecting all middle clusters, *P*^*g*^ is the final updated *P*^*s*^.

We use a standard HAC tree to perform MIRAC when the number of samples is larger than ten thousand, which usually results in decreased but still acceptable performances (**Figures 2**). Although the optimal leaf ordering of the HAC tree is an important assumption, its computation takes too long when the number of samples is large. For example, it takes more than a week to compute the optimal leaf ordering of a dataset with about 68,000 samples. However, the time complexity of computing a good HAC linear ordering could be significantly reduced by implementing other ordering techniques (Aydin, Bateni, and Mirrokni 2016).

**Figure 2.**
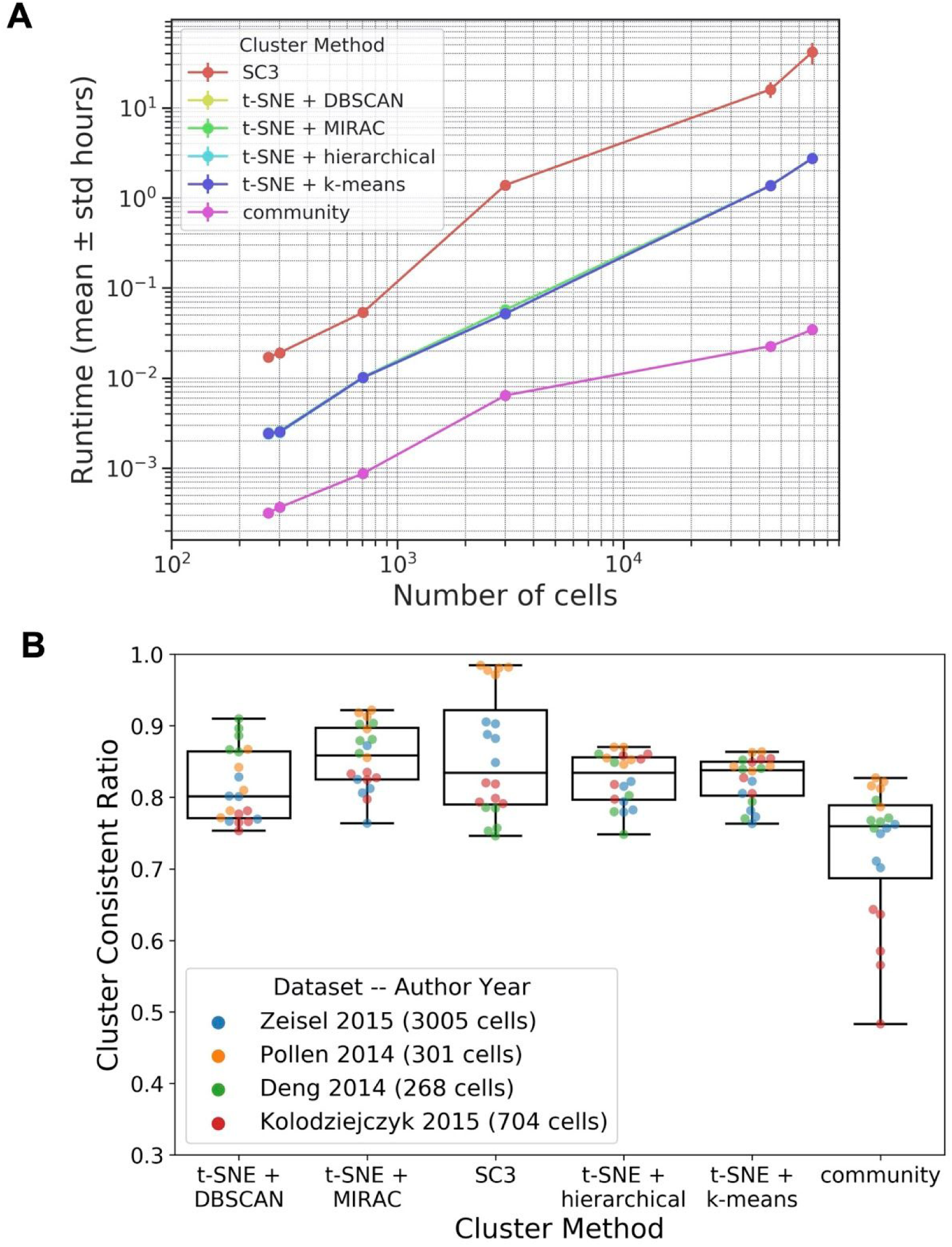
Clustering method benchmarks on experimental datasets. (**A**) Runtimes. (**B**) CCRs on different datasets, with different points of each dataset representing different numbers of clusters. For each dataset, the numbers of clusters are the same across all compared clustering methods.

#### 2.2.2 Analysis of complexity

The time complexity of MIRAC is *O*(*n*^4^ + *mn*^2^), where *n, m* ≥ 1. The time complexity of computing the HAC tree with optimal leaf ordering is *O*(*n*^4^) (Bar-Joseph, Gifford, and Jaakkola 2001). The time complexity of finding *P*^*g*^ is *O*(*mn*^2^), which is briefly analyzed as following.

The time complexity is *O*(*mn*) for computing the code length *L*(*X*_*n*×*m*_, *P*). In computing *L*(*X*_*n*×*m*_, *P*), we compute the code lengths of each cluster and their labels. Since it takes *O*(*n*) time to compute the code length of *n* observations with a single feature, computing the code length of all clusters and features individually takes *O*(*mn*) time, and computing the code length of *n* labels take *O*(*n*) time.

Similarly, the time complexity is *O*(*n*_1_*m* + *n*_2_*m*) for computing the code length of 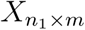 encoded by a model fitted with 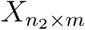. Fitting a model on 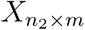 takes *O*(*n*_2_*m*) amount of time. Using the model to encode 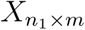 takes *O*(*n*_1_*m*) amount of time.

The time complexity of searching for *P*^*g*^ is *O*(*mn*^2^). In the worst case scenario, |*P*^*s*^| = *n*, and the inspecting cluster is always merged with the left hand side cluster in step 4. However, the impact of 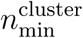 and step 3 on the time complexity is not straightforward, so we divide step 3 code length computation into three parts and analyze the upper bound of them individually. The divided three parts are the left cluster, inspecting cluster, and right minimax cluster. Because we keep increasing the size of the left cluster, the upper bound of computing the left cluster code length throughout the execution is *O*(*mn*^2^). The maximum size of the inspecting cluster is 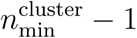, so the upper bound of computing the inspecting cluster code length is also *O*(*mn*^2^), when every middle singleton is evaluated 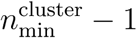 times before merging with the left. The maximum size of the right minimax cluster is 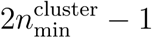,so the upper bound of evaluating right minimax cluster is also *O*(*mn*^2^), when every increment of the left cluster takes 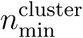 times of evaluating the right minimax cluster. Summarizing these three parts, the overall upper bound is *O*(*mn*^2^).

The space complexity of MIRAC is *O*(*nm* + *n*^2^), which could be decomposed into the following three parts. The space complexity of storing *n* × *m* the data matrix *X* is *O*(*nm*). The space complexity of storing the HAC tree with optimal leaf ordering is *O*(*n*). The space complexity of storing the pairwise distance matrix is *O*(*n*^2^).

#### 2.2.3 Extension with community detection

We extended MIRAC with community detection to improve scalability. We apply MIRAC on a relatively large number of detected single-cell communities to identify final clusters. We used the Leiden algorithm for community detection on KNN graphs (Traag, Waltman, and van Eck 2019). We also provide a KNN graph construction method that supports approximate nearest neighbor (ANN) search using a Hierarchical Navigable Small World (HNSW) graph (Malkov and Yashunin 2016).

### 2.3 Cluster separating genes identification

We use XGBoost (Chen and Guestrin 2016), a scalable and sparsity-aware boosted tree system, to identify genes that are able to separate a specific set of clusters. This method is designed for data exploration after applying any one of a number of various statistical approaches that have been developed to identify differentially expressed genes (Vallejos, Marioni, and Richardson 2015; Kharchenko, Silberstein, and Scadden 2014; Qiu et al. 2017; Soneson and Robinson 2018; Korthauer et al. 2016), in order to quickly identify genes or sets of genes that are able to separate an arbitrary set of clusters for further inspection.

Rather than providing a meticulous p-value for each gene among the compared clusters, we rank the genes by their importances on separating the clusters under comparison. The importance of a gene in the trained classifier is the number of times it has been used as an inner node of the decision trees. We use cross validation to train an XGBoost classifier on the compared clusters, and the classifier is essentially a bag of decision trees (Chen and Guestrin 2016). In order to alleviate the influences of stochasticity on interpretation, we use bootstrap with feature shuffling to better estimate the importance of genes in separating the compared clusters. The obtained list of important genes could be further explored by inspecting transcription level fold changes and decision tree structures.

Comparing to NSForest (Aevermann et al. 2018), a method based on random forest (Breiman 2001; Pedregosa et al. 2011) to identify a parsimonious set of cluster separating genes from scRNA-seq data, our method identifies all possible cluster separating genes using gradient boosting. Practically, NSForest version 1.3 is distributed on GitHub as a Python script without encapsulation or testing (checked on Oct 11, 2018), but our method is distributed through the Python Package Index, comprehensively tested, and easy to use through the user friendly API. With regard to scalability, our method uses a scalable implementation of gradient boosting algorithm (Chen and Guestrin 2016), whereas NSForest uses the implementation of random forest in scikit-learn (Pedregosa et al. 2011).

### 2.4 K-nearest neighbor methods

A k-nearest neighbor analytical strategy exploits the similarity between data points for classification, regression, and imputation (Cover and Hart 2006; Bezdek, Chuah, and Leep 1986). In k-nearest neighbor methods, samples are considered as points in a space with each dimension representing a measured property, which is often referred to as a feature, of the samples. The similarity between samples can be evaluated with various distance metrics, which is extensively reviewed by Bellet *et al*. (Bellet, Habrard, and Sebban 2013). Generally, a distance metric is a function that takes two samples and output a numeric value to represent the distance between the two samples in their feature space. For example, the Euclidean distance metric is the following function, 

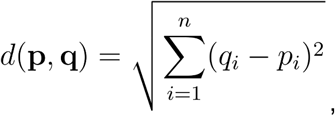

where the **P** and **q** are two samples, and and *q*_*i*_ are *p*_*i*_ the data values on their *i*th dimension. The k-nearest neighbors of a sample are the k number of other samples that have the smallest distances from the sample. The k-nearest neighbors of a sample are informative, due to their similarity, to determine the category of the sample in classification, the relevant continuous property in regression, and the missing values in imputation.

We developed two methods based on the KNN algorithm to facilitate the exploration of scRNA-seq datasets. With relative large number of cells profiled in each scRNA-seq experiment, we assume that each one of the non-rare cells is similar to at least *k* other cells in their transcriptomic profiles. With this assumption, we impute gene dropouts and detect rare transcriptomic profiles. In the scedar implementation, we also support approximate nearest neighbor (ANN) search using Hierarchical Navigable Small World (HNSW) graph (Malkov and Yashunin 2016), which greatly improves the scalability of KNN graph construction.

#### 2.4.1 Impute gene dropouts

In an scRNA-seq experiment, if a truly expressed gene is not detected in a cell, the gene is considered a “dropout”, and such events are called gene dropouts (Kharchenko, Silberstein, and Scadden 2014). Gene dropouts may be caused by biological and technical reasons (Kharchenko, Silberstein, and Scadden 2014; Risso et al. 2018; Pierson and Yau 2015). The rationale behind the possible causes of biological dropouts mainly involves transcriptional bursting (Suter et al. 2011; Tantale et al. 2016) and RNA degradation. With regard to technical dropouts, the main concerns are the relatively small number of RNA transcripts of a gene, amplification efficiency, and batch effect (Zappia, Phipson, and Oshlack 2017).

We exploit the transcriptomic profiles of the k-nearest neighbors of a cell to impute the gene dropouts in the cell (**Algorithm S1**). The algorithm could take multiple iterations, so that the dropped-out genes that are expressed in all k-nearest neighbors could be imputed at first, and the ones that are expressed in most but not all of k-nearest neighbors could be imputed in the following iterations.

#### 2.4.2 Detecting rare transcriptomic profiles

We mark transcriptomic profiles as rare if they are distinct from their k-nearest neighbors, according to the pairwise similarity between cells (**Algorithm S2**). The algorithm could take multiple iterations, so that the most distinct transcriptomic profiles could be marked at first and less distinct ones in the following iterations.

This method is provided mainly to facilitate detailed inspection of rare transcriptomic profiles rather than removing outliers from the data. Because rare transcriptomic profiles may have various biological and technical causes, samples and features in a dataset should only be removed after extensive exploratory data analysis and rigorous reasoning with domain specific knowledge. Closely comparing rare transcriptomic profiles with their nearest neighbors may also yield insights into their biological differences, which may further facilitate the identification of rare cell types and states.

### 2.5 Benchmark

We benchmarked the clustering and KNN performances of scedar on simulated and experimental scRNA-seq datasets. We obtained previously published experimental scRNA-seq datasets (**Table 1**). We also generated 50 simulated scRNA-seq read count datasets with Splatter (Zappia, Phipson, and Oshlack 2017). The simulation parameters are estimated by Splatter according to a Drop-seq dataset (Macosko et al. 2015) and a 10x Genomics GemCode dataset (Zheng et al. 2017). Within each simulated dataset, the cells have 8 clusters taking approximately the following proportions: ⟨0.3, 0.2, 0.15, 0.15, 0.05, 0.05, 0.05, 0.05⟩, with a gene dropout rate around 5%.

**Table 1.** Real scRNA-seq datasets for benchmark

| Publication | # cells | # genes | Organism | Tissue | Protocol | Raw Data Type |
| --- | --- | --- | --- | --- | --- | --- |
| Deng *et al.* (2014) | 268 | 22,431 | Mus Musculus | Embryo | Smart-Seq/Smart-Seq2 | Read count |
| Pollen *et al.* (2014) | 301 | 23,730 | Homo sapiens | Dermal, blood, pluripotent and neural | SMARTer | TPM |
| Kolodziejczyk *et al.* (2015) | 704 | 38,653 | Mus Musculus | Embryonic stem cell | SMARTer | Read count |
| Zeisel *et al.* (2015) | 3005 | 19,972 | Mus Musculus | Brain | STRT-Seq UMI | Read count |
| Macosko *et al.* (2015) | 44,808 | 23,288 | Mus Musculus | Retina | Drop-seq | Read count |
| Zheng *et al.* (2015) | 68,579 | 33,694 | Homo sapiens | Peripheral blood mononuclear cell | GemCode | Read count |
| Han *et al.* (2018) | 405,191 | 39,855 | Mus Musculus | Various tissues | Microwell-Seq | Read count |
| Cao *et al.* (2019) | 2,058,652 | 26,183 | Mus Musculus | Embryo | sci-RNA-seq3 | Read count |

Source: https://hemberg-lab.github.io/scRNA.seq.datasets/. TPM represents Transcripts Per Million in the raw data type column.

We performed all benchmark analyses on a high-performance computing cluster, of which the computing resources are strictly managed by Univa Grid Engine. The cluster nodes have CPUs of Intel Xeon E5-2680 v3 or Intel Xeon E7-8880 v3. The memory sizes are either 128GB, 256GB, or 1TB. When scheduling analytical jobs for benchmarking, we make sure that the number of cores and allocated memory are enough for the program.

#### 2.5.1 Clustering

The clustering accuracy and stability of MIRAC were benchmarked together with several other clustering methods on experimental scRNA-seq datasets listed in **Table 1**.

The following clustering methods are directly applied on the original data without preprocessing.

- MIRAC on 2D t-SNE projection.
- Single-cell consensus clustering (SC3) version 1.7.7 (Kiselev et al. 2017). SC3 is selected for comparison because it was extensively compared with other methods on different experimental datasets.
- K-means clustering on 2D t-SNE projection.
- Hierarchical agglomerative clustering on 2D t-SNE projection.
- Density-based spatial clustering of applications with noise (DBSCAN) (Ester et al. 1996) on 2D t-SNE projection.
- Community clustering on KNN graph constructed using the Euclidean distances of 100 principal components. We used the Leiden algorithm for community detection (Traag, Waltman, and van Eck 2019).

Although MIRAC could be directly applied on the expression matrix, dimensionality reduction is able to improve the performance of similarity and density based clustering methods when the number of features is high (Aggarwal, Hinneburg, and Keim 2001). The mathematical influences of the high number of features are briefly described in the previous sections.

Although t-SNE projections are stochastic and influenced by the perplexity parameter, t-SNE is proved to be able to recover well-separated clusters (Linderman and Steinerberger 2019)and t-SNE has been extensively used as dimensionality reduction method for scRNA-seq data (Macosko et al. 2015; Cao et al. 2017). Thus we chose to use t-SNE for r demonstration in this report, but we note that scedar also supports the use of PCA and UMAP for MIRAC clustering, which can be applied just as easily as the t-SNE method.

When benchmarking for accuracy, we cluster the cells in each experimental dataset using the compared clustering methods with a grid of parameters. We use the maximum similarities between the clustering results and the cell types from the publications in order to compare the accuracy of different clustering methods. Although taking the maximum increases the chance of overfitting, it resembles the procedure of clustering analysis in practice.

We also recorded the runtime of clustering methods on different datasets. In **Figure 2A**, SC3 was performed with 20 cores, and community clustering is performed with 60 cores. Other clustering methods were performed with a single core. When running MIRAC, we did not require the hierarchical tree to be optimal. In **Figure S8C**, the community-detection-extended MIRAC clustering method was performed with 25 CPU cores, and other methods were performed with 60 CPU cores.

We also benchmarked the influences of t-SNE random states on the clustering results (Figure S1). T-SNE embeddings generated with different random states may be distinct from each other, although current mathematical results have shown that clustering using t-SNE embeddings is able to recover well separated clusters (Linderman and Steinerberger 2019). We characterized the stability to random states of clustering methods by running them on each experimental dataset with the same parameters but ten different random states. The similarity of clustering results between different random states are used to compare the stability of different clustering methods. The stability of community clustering is not characterized, because it is based on PCA.

#### 2.5.2 Cluster similarity metrics

We use two cluster similarity metrics, cluster consistent ratio (CCR) and adjusted Rand index (ARI) (Hubert and Arabie 1985) for measuring the accuracy and stability of clustering methods respectively. When we have a coarse reference partition *P*_*r*_ and a finer clustering partition *P*_*c*_, the CCR is computed as the ratio of pairs within each cluster of *P*_*c*_ that are also in the same cluster of *P*_*r*_, with the number of clusters kept the same across compared methods. The ARI is computed with the Python package scikit-learn (Pedregosa et al. 2011) using the mathematical formula given by Hubert and Arabie (Hubert and Arabie 1985).

The reference partitions *P*_*r*_ of real datasets are obtained from their original publications (**Table S1**). The clusters in Deng *et al*. dataset (Deng et al. 2014) are experimentally determined by manual single cell isolation. The clusters in Pollen *et al.* (Pollen et al. 2014) are experimentally determined by the cell line or tissue type of the isolated single cells. The clusters in Kolodziejczyk *et al*. (Kolodziejczyk et al. 2015) are determined by the embryonic stem cell culturing conditions and batches. The clusters in Zeisel *et al.* (Zeisel et al. 2015) are determined by the BackSPIN clustering method (Zeisel et al. 2015) and further inspected with domain specific knowledge.

We choose CCR to measure clustering accuracy rather than ARI, because ARI greatly penalizes the split of a large cluster in *P*_*r*_ into multiple smaller ones in *P*_*c*_. This behavior of ARI could prevent the evaluation of transcriptomic variabilities within a large group of cells in clustering analysis, which is an important goal of scRNA-seq experiments that cannot be easily achieved by bulk RNA-seq experiments. In addition, we also provide a method in scedar to easily merge multiple clusters together, in case the users found the sub-types of a cell type are very similar to each other.

However, we used ARI to measure clustering stability rather than CCR, because the differences between *P*_*r*_ and *P*_*c*_ are completely caused by different random states, hence splitting a cluster in *P*_*r*_ should be penalized.

MDL is not used as a cluster similarity metric, even though MDL is used in MIRAC to guide the process of finding good partitions. MDL is only used to guide and regularize the merging process of local adjacent sub-clusters by evaluating their similarity, but it is not used as an objective to be optimized globally. Using MDL as a benchmarking metric would introduce a bias as among the various methods, MIRAC is the only method using MDL. Moreover, interpreting relative global MDL differences is not straightforward as it is not only affected by cluster labels, but also transcription levels.

#### 2.5.3 Detection of rare transcriptomic profiles

We visualize real datasets before and after removing rare transcriptomic profiles in t-SNE projection and pairwise distance heatmaps, without quantitative evaluations like receiver operating characteristic (ROC) curve, since rare transcriptomic profiles are not well defined with a large number of genes (Aggarwal, Hinneburg, and Keim 2001). Approaches to identify rare data points in low-dimensional spaces (≤ 3) do not scale well to higher dimensions due to the exponentially decreased density of data points in space and increased instability of distances, which is elaborated in the previous section about mathematical theories on high-dimensional data analysis. In **Figure S8B**, KNN rare transcriptomic profile detection is performed with 25 CPU cores.

It is important to note that rare transcriptomic profiles are detected to facilitate detailed inspection rather than the removal of them from the data. The visualizations are only used to illustrate the capability of the KNN method for detecting rare transcriptomic profiles.

#### 2.5.4 Gene dropout imputation

We simulate gene dropouts using Splatter (Zappia, Phipson, and Oshlack 2017) to obtain a dropout rate around 5%. Then, we benchmark the performance of imputing gene dropout as two parts: detection and smoothing. On simulated data, because the true gene dropouts are known, we use a ROC curve and mean squared errors (MSEs) to characterize the performance of gene dropout detection and smoothing respectively. In **Figure 3A, S7A, and S8A**, all imputation methods were executed with a single core. On real data, we visualize the cells in 2D t-SNE space before and after imputation. The compared methods are KNN gene dropout imputation (KNNGDI) version 0.1.5, SAVER version 1.0.0 (Huang et al. 2018), MAGIC version 1.1.0 (van Dijk et al. 2018), and scImpute version 0.0.6 (Li and Li 2017).

**Figure 3.**
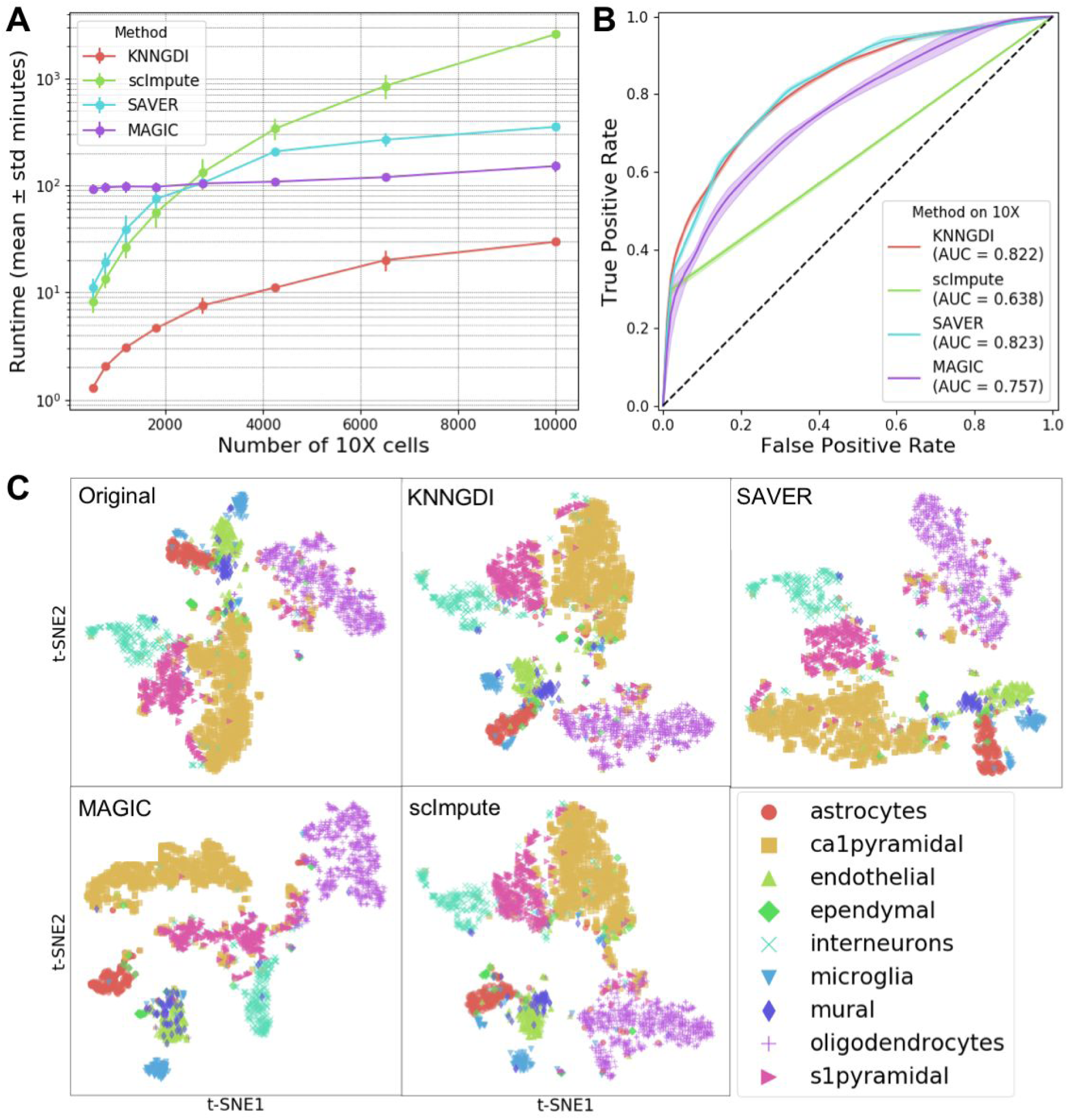
Gene dropout imputation method benchmarks. (**A**) Runtimes on 40 simulated 10x Genomics datasets. (**B**) ROC curves (± standard deviation) of dropout detection on the simulated 10x Genomics datasets. (**C**) t-SNE scatter plots of the Zeisel *et al*. (Zeisel et al. 2015) dataset after gene dropout imputations.

#### 2.5.5 Method speed-up by parallel processing

We characterized the analysis speed-up by parallel processing on different datasets (**Figure S9**). KNN dropout imputation is not evaluated on larger datasets, because the speed limiting step has not yet been parallized. In addition to parallelizing the computation in the implemented methods, we also provide an easy-to-use utility function (scedar.utils.parmap) to parallely run analytical methods with different parameters, which would significantly facilitate exploratory parameter search.

## 3 Results

### 3.1 Basic workflow of scedar

We illustrate the basic workflow of using scedar for scRNA-seq exploratory data analysis with the dataset published by Zeisel *et al.* (Zeisel et al. 2015) (**Figure 1**). The dataset contains the RNA unique molecule identifier (UMI) counts of 19,972 genes in 3005 cells from the mouse brain. We selected this dataset for demonstration because it could be clearly visualized in small graphs, although our package is capable of analyzing much larger datasets with over 2 million cells (**Figures S2, S3, S4** and **S5**).

In **Figure 1**, each box represents a data analysis step, and they are consecutively executed according to the arrow. The purpose and runtime are listed in the upper ribbon. The code that is essential to the step is listed in the box, and their results are also shown.

The preparation step imports required packages and loads the data. The class SampleDistanceMatrix is one of the core data structures in the package that is used to store the read counts and pairwise distances of an scRNA-seq dataset. Because the pairwise distance computation is delayed until necessary, i.e. lazily loaded, this step only takes 12 seconds. We use cosine distance rather than correlation distance to greatly speed up the computation, since we implemented the computation procedure of pairwise cosine distances completely with numpy linear algebra operations with OpenBLAS backend (Xianyi, Qian, and Yunquan 2012; Wang et al. 2013).

The t-Distributed Stochastic Neighbor Embedding (t-SNE) scatter plot and KNN graph are used to explore the dataset. The cell type labels published by Zeisel *et al.* (Zeisel et al. 2015) are truncated to fit in the space. We also provide methods to visualize arbitrary statistics, e.g. number of expressed genes, of individual cells as color gradient. The layouts of cells in t-SNE and KNN graph are similar to each other. Although KNN graph is faster than t-SNE, the runtime for t-SNE could be reduced by optimizing its parameters, e.g. lowering the number of iterations.

The MIRAC step clusters the cells and visualizes them with t-SNE scatter plot and pairwise distance matrix heatmap. The heatmap generation procedure in scedar is optimized for large-scale datasets, which is able to generate a heatmap with tens of thousands of columns and rows in a few minutes. Users could also generate heatmaps for the read count matrix to directly inspect the sparsity of datasets (**Figures S2B, S3B, S4B** and **S5B**).

The last step identifies cluster separating genes with XGBoost (Chen and Guestrin 2016). Users could choose an arbitrary set of clusters to compare, and the genes are ranked by their importance in separating the clusters. Then, the read counts of a gene across clustered labels could easily be visualized by t-SNE scatter plot.

### 3.2 Performance of MIRAC clustering

We benchmarked several clustering methods on the datasets listed in **Table 1** (**Figure 2**). Each dataset is clustered multiple times with each clustering method to obtain different numbers of clusters (**Table S1**).

The t-SNE based clustering methods are faster than SC3 (**Figure 2A**). Also, the t-SNE based clustering methods have similar runtimes (**Figure 2A)**, since the time limiting step is the computation of t-SNE projection, when optimal hierarchical clustering ordering is not required in MIRAC. When the optimal ordering is required, the time limiting step is the computation of optimal ordering (**Figure 1**). With regard to practical usage, we recommend the users to explore different parameters of MIRAC without optimal ordering. Once appropriate parameters have been identified, users can perform MIRAC with optimal ordering. However, for datasets with more than 10,000 cells, we recommend that users perform MIRAC without optimal ordering (**Figure S2C and S3C**). For datasets with more than 100,000 cells, we recommend that users perform community-detection-extended MIRAC without optimal ordering (**Figure S8C**), which takes for instance 2885.5 ± 176.0 (mean ± standard deviation) seconds on the mouse organogenesis cell atlas (MOCA) dataset with 2,058,652 single cells using 25 CPU cores (**Figure S8C**) (Cao et al. 2019). We also performed clustering with the community clustering methods implemented in scedar, Seurat (Butler et al. 2018) and Scanpy (Wolf, Angerer, and Theis 2018) on relatively large scRNA-seq datasets with more than 10,000 cells (**Figure S8C**). The runtime of community clustering methods Seurat and Scanpy is comparable to community clustering and community-detection-extended MIRAC (**Figure S8C**), except that Scanpy and Seurat were not able to cluster the MOCA dataset that contain 2,058,652 single cells on a server with 1TB memory due to a mandatory conversion of the sparse read count matrix into dense matrix.

The cluster consistent ratios (CCRs) of t-SNE based clustering methods are comparable to SC3 (**Figure 2B**). The PCA based community clustering results showed relatively lower CCRs than other clustering methods (**Figure 2B**), which might due to the non-optimal representation of similarities between high-dimensional single cell read counts in the linear PCA space, comparing to t-SNE embeddings that are optimized to capture the similarities between high-dimensional single cell read counts. The representative MIRAC clustering results of Zheng *et al.* (Zheng et al. 2017) and Macosko *et al.* (Macosko et al. 2015) datasets are visualized with t-SNE scatter plots and pairwise distance matrix heatmaps (**Figures S2C** and **S3C**). For smaller datasets, the representative MIRAC results are shown in **Figure S6**. Although t-SNE projections obtained with different random states are distinct from each other, the consistency of clustering results is comparable to SC3 (**Figure S1**).

### 3.3 Performance of imputing gene dropouts

We benchmarked several gene dropout imputation methods on the simulated 10x Genomics (**Figure 3**) and Drop-seq (**Figure S7**) datasets. K-nearest neighbor gene dropout imputation (KNNGDI) is faster than other compared methods on relatively small datasets (**Figure 3A**) and is capable of handling scRNA-seq datasets with over 10,000 cells (**Figure S8A**). The runtime of KNNDGI on 2,058,652 cells is 71.56 ± 1.71 (mean ± standard deviation) hours.

The performance of KNNGDI on detecting gene dropouts is comparable to SAVER and better than scImpute and MAGIC. The ROC curve of scImpute sharply turns around TPR = 0.25 (**Figure 3B** and **S7B**) because its threshold parameters, determining whether a zero entry is a dropout or not, are not sensitive enough to achieve any higher TPRs. Although the AUCs of KNNGDI and SAVER are higher than scImpute and MAGIC, they all have comparable performances when FPRs are lower than 0.05 (**Figure 3B**), except that MAGIC has worse performance on the simulated Drop-seq datasets that have higher sparsity than the Zeisel *et al.* (Zeisel et al. 2015) dataset (**Figure S7B**).

The MSEs of KNNGDI on correcting gene dropouts are comparable to SAVER and smaller than scImpute and MAGIC (**Figure S7C** and **S7D**). However, these methods all have MSEs many times higher than the MSEs of the observed counts to the true counts, which implies that these methods all introduced many times more reads than the true dropout reads. Despite different levels of MSEs, the t-SNE embeddings of the imputed read counts are consistent with the t-SNE embeddings of original read counts (**Figure 3C, S2E, S3E, S4E and S5E**).

### 3.4 Performance of detecting rare transcriptomic profiles

We detected rare transcriptomic profiles in datasets listed in **Table 1** with the KNN method (**Figure 4, S2F, S3F, S4F, S5F** and **S6**). Although the detection method has many limitations, rare transcriptomic profiles tend to be points on the t-SNE scatter plots either far away from the majority of their same types or along the edges of a group of agglomerated points. On the pairwise distance matrix heatmap, rare transcriptomic profiles tend to be small chunks that are distinct from their neighbors, and the heatmap becomes smoother after removing the rare transcriptomic profiles. The detected rare transcriptomic profiles could be further inspected as potential rare cell types and states by comparing with their nearest neighbors. The KNN rare transcriptomic profile detection method is also capable of handling scRNA-seq datasets with over 10,000 cells (**Figure S8B**), with a runtime of 22.14 ± 0.87 (mean ± standard deviation) minutes on 2,058,652 cells.

**Figure 4.**
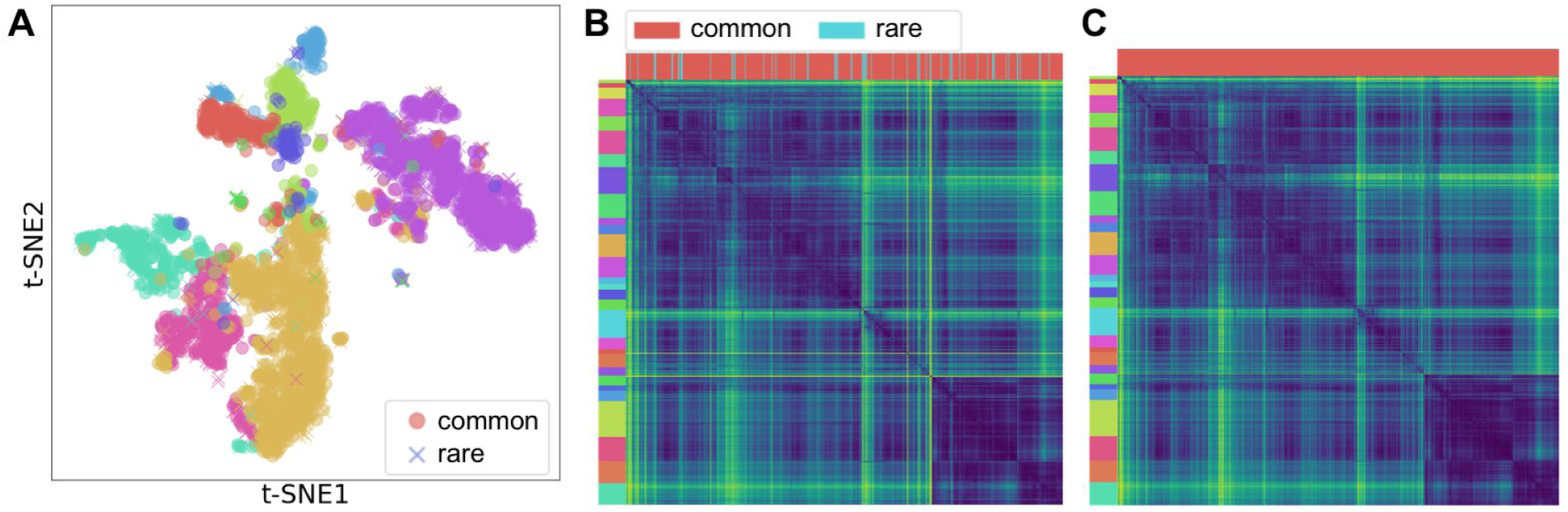
KNN rare transcriptomic profile detection on the Zeisel *et al*. (Zeisel et al. 2015) dataset. (**A**) t-SNE scatter plot with colors labeling cell types and markers labeling common or rare transcriptomic profiles. 9.3% cells are marked as rare. (**B**) Pairwise cosine distance heatmap with left strip as MIRAC labels and upper strip as common or rare transcriptomic profiles labels. (**C**) Pairwise cosine distance heatmap with rare transcriptomic profiles removed.

### 3.5 Identification of cluster separating genes

We used scedar to identify the genes distinguishing the MIRAC cluster 1, 15, and 22 of the Zeisel *et al*. (Zeisel et al. 2015) dataset (**Figure 5** and **Table S2**). In the original publication, the upper to lower MIRAC clusters 22, 1, and 15 in the t-SNE scatter plot are assigned to microglia, endothelial cells, and astrocytes respectively (**Figure 3C**). We choose these three clusters to inspect one of the discrepancies between cell types and MIRAC clustering results, where the small isolated upper part of the MIRAC cluster 15 is assigned to microglia instead of endothelial cells in the original publication.

**Figure 5.**
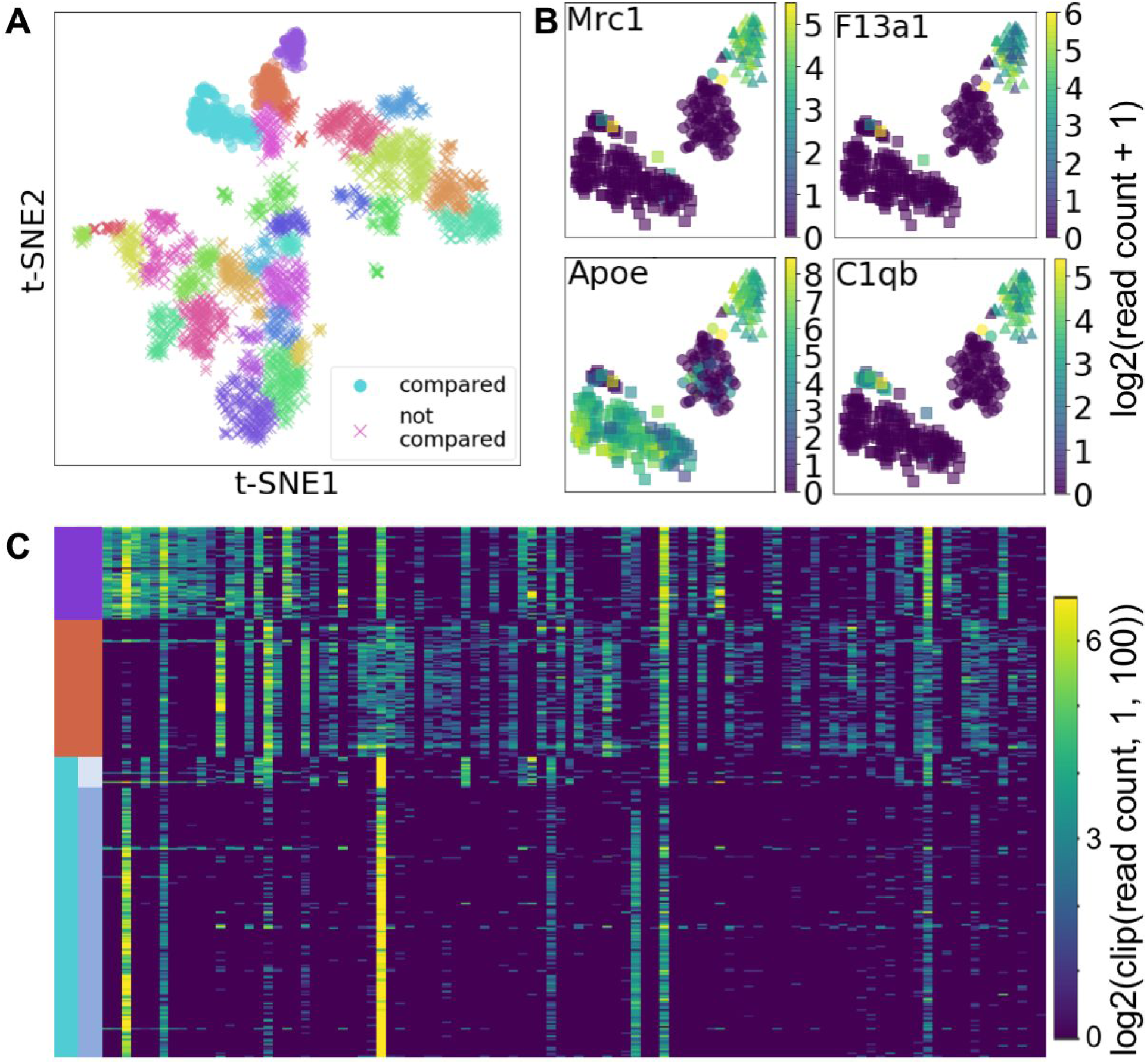
Identified genes separating the MIRAC clusters 1, 15, and 22 of the Zeisel *et al*. (Zeisel et al. 2015) dataset. (**A**) t-SNE scatter plot with color as MIRAC cluster labels and marker shape as compared or not compared. (**B**) t-SNE scatter plots of the compared clusters with color as log_2_(read count + 1) of the corresponding gene and marker shape as MIRAC clusters. (**C**) Transcription level heatmap of the top 100 important cluster separating genes in the compared cells, with rows as cells ordered by cluster labels and columns as genes ordered by importance. The color gradient is log_2_(clip(read count, 1, 100)), where the clip(read count, 1, 100) function changes any read count below 1 to 1 and above 100 to 100, in order to better compare genes at different transcription levels.

The smaller upper isolated part of MIRAC cluster 15 might be a distinct cell sub-type of microglia. Although it expresses a microglia marker gene C1qb (**Figure 5B**) (Beutner et al. 2013), it does not express Mrc1 or Apoe in the same pattern as the MIRAC cluster 22 (**Figure 5B** and **5C**). According to the transcription levels of the cluster separating genes (**Figure 5C**), the smaller upper isolated part of MIRAC cluster 15, which is located at the top of the cluster 15 rows, has some genes expressed in the same pattern as the microglia, but it also has some other genes expressed distinctly from the microglia.

## 4 Discussion

Comprehensive profiling of the transcriptomes of individual cells within an organism or tissue by scRNA-seq is able to facilitate the systematic study of physiological or pathological states (Villani et al. 2017; Baslan and Hicks 2017; Cao et al. 2017). Previous scRNA-seq experiments identified novel cell types or states (Villani et al. 2017), obtained insights into the regulation of differentiation process (Cao et al. 2017; Rizvi et al. 2017; Cao et al. 2019; Schiebinger et al. 2019), and inferred molecular mechanisms of tissue functions (Zeisel et al. 2015; Lake et al. 2018; Haber et al. 2017).

The biological results of scRNA-seq experiments are obtained from extensive data analyses, which could take more time than doing the experiments. As the size of scRNA-seq datasets increases rapidly (Svensson, Vento-Tormo, and Teichmann 2018; Rozenblatt-Rosen et al. 2017), computational methods also need to be scalable by exploiting hardware and software techniques for big data analytics, in order to complete an analysis within a reasonable amount of time.

Scedar is able to facilitate gene dropout imputation, rare transcriptomic profile detection, clustering, and identification of cluster separating genes for scRNA-seq data by exploiting scalable system design patterns and high-performance computing architectures. We parallelized time-consuming computational procedures by multi-processing without excessive copies of shared data in memory. We also decompose the whole analytical procedure into multiple steps, so that certain steps could be specifically optimized without repeating others. In addition, intermediate results, such as pairwise distances and t-SNE projections, are lazy loaded and cached in a unified data structure to speed up analytical routines, prevent repeated computations, and alleviate the burden on users to keep track of all intermediate results.

Comparing to other computational tools that were developed or updated for large scale scRNA-seq data analysis, like PAGODA2 (Fan et al. 2016), Seurat v2.0 (Butler et al. 2018), and Scanpy (Wolf, Angerer, and Theis 2018), scedar distinguishes itself with an additional research focus on developing new analytical methods, including those based on machine learning. In scedar, we adapted KNN, a typical machine learning algorithm, to impute gene dropouts and detect rare transcriptomic profiles. MDL principle, an important concept in computational learning, is applied to cluster single cells (Hansen and Yu 2001). A scalable and sparsity-aware gradient boosted tree system XGBoost (Chen and Guestrin 2016), which implements a typical machine learning algorithm, is used to identify genes that are able to distinguish different clusters. In addition, these machine learning methods are able to exploit modern high-performance computing architecture, which improves the scalability of the package.

In scedar, we developed a clustering algorithm, MIRAC, for scRNA-seq data. MIRAC clusters observations in three steps: 1) build a tree by hierarchical clustering, 2) divide the tree into sub-trees, and 3) merge similar sub-trees into individual clusters. This clustering strategy adapts the basic ideas of BIRCH (Zhang, Ramakrishnan, and Livny 1996) and BackSPIN (Zeisel et al. 2015). Instead of building a balanced tree structure, like the clustering feature tree in BIRCH for further partitioning, MIRAC divides the tree structure built by hierarchical clustering optionally in a balanced manner (**Figure S10** and Supplementary Method section 1.4), which simplifies the clustering procedure. In BackSPIN, a sorted pairwise correlation matrix is recursively bi-partitioned into clusters according to criteria based on the normalized sums of correlation coefficients. In contrast, MIRAC iteratively merges the relatively small sub-clusters according to criteria based on the MDL principle, which increases the robustness for determining whether two groups of observations should be put in the same cluster or not. Especially, when the number of observations is large within a group, finding an optimal bi-partition is not straightforward, since there may be multiple distinct sub-groups. Although the performance of MIRAC under certain metrics is comparable to other clustering algorithms, it is able to provide distinct clusters that are sensitive to local structures, which could be used as alternative perspectives to interpret the source of heterogeneity within the dataset.

There are still many possible improvements on scedar. To improve the scalability of MIRAC, we could provide more efficient methods to obtain the optimal leaf ordering using linear embedding techniques (Aydin, Bateni, and Mirrokni 2016). To improve the scalability of the backend data structure, we could extend it with distributed analytic systems such as Apache Spark (Zaharia et al. 2016). To visualize the differences between different clusters, we could plot the fold changes of gene transcription levels on the pathway maps in the KEGG (Kyoto Encyclopedia of Genes and Genomes) database (Kanehisa et al. 2016).

## Supporting information

Supplementary-scedar-2020

## 5 Data availability

In scedar benchmark, the real scRNA-seq datasets can be obtained from the following websites:

- https://hemberg-lab.github.io/scRNA.seq.datasets
- https://support.10xgenomics.com/single-cell-gene-expression/datasets
- https://figshare.com/articles/MCA_DGE_Data/5435866
- https://oncoscape.v3.sttrcancer.org/atlas.gs.washington.edu.mouse.rna/downloads

The simulated datasets can be generated using scripts at https://github.com/logstar/scedar/tree/master/docs/r_scripts.

## 6 Code Availability

The scedar package is distributed through Pypi https://pypi.org/project/scedar

The code and other resources are available in Github at https://github.com/logstar/scedar

## 7 Supplementary Text

We have provided a Supplementary file with text, algorithms and figures referenced in this work.

