## Supplementary-scedar-2020 for "Scedar: a scalable Python package for single-cell RNA-seq exploratory data analysis"

Yuanchao Zhang[1,2], Man S. Kim[1], Erin R. Reichenberger[1], Ben Stear[1], and Deanne M. Taylor[1,3]
1. Department of Biomedical and Health Informatics, The Children's Hospital of Philadelphia, Philadelphia, PA 19041, USA
2. Department of Genetics, Rutgers University, Piscataway, NJ 08854, USA
3. Department of Pediatrics, Perelman School of Medicine, University of Pennsylvania, Philadelphia, Pennsylvania, 19104, USA

### 1 Supplementary methods

#### 1.1 scedar package development

Scedar is built upon various high-performance scientific computing and visualization packages. Scedar is also extensively benchmarked and tested by unit testing, with comprehensive coverage on statements and branches.

At the time of publication, scedar uses the following packages and versions:

- numpy version 1.18.1 (Oliphant 2006) for matrix representation and operations.
- scipy version 1.4.1 (Virtanen et al. 2018) for fast Gaussian kernel density estimation, hierarchical clustering and sparse matrix.
- matplotlib version 3.1.0 (Hunter 2007) and seaborn (Waskom et al. 2017) for visualization.
- pandas version 0.25.3 (McKinney and Others 2010) for data frame representation.
- scikit-learn version 0.21.0 (Pedregosa et al. 2011) for parallel computation of pairwise distances, k-nearest neighbor (KNN) data structure, PCA and t-SNE.
- XGBoost version 0.90 (Chen and Guestrin 2016) for scalable gradient boosting tree.
- networkx version 2.4 (Hagberg, Swart, and S Chult 2008) for graph data structure and visualization.
- ForceAtlas2 version 0.3.5 (Jacomy et al. 2014) for scalable force-directed graph layout.
- nmslib version 2.0.5 (Malkov and Yashunin 2016) for approximate nearest neighbor search using Hierarchical Navigable Small World Graphs.
- Leidenalg version 0.7 (Traag, Waltman, and van Eck 2019) for community detection.
- umap-learn version 0.3.10 (McInnes, Healy, and Melville 2018; McInnes et al. 2018) for computing UMAP embeddings.

- python-igraph version 0.7.1.post6 (Csardi and Nepusz 2006) for constructing KNN graph data structure.

We use the Python package pytest version 5.2.0 (Krekel et al. 2004) as the unit testing framework to ensure that scedar has expected behaviors, even after major coding changes. We tested each member of the package with multiple testing environments, in order to make sure that all statements and branches are executed in the tests, i.e. comprehensive code coverage. The code coverage was measured by the Python package coverage (v 5.03, https://coverage.readthedocs.io/en/coverage-5.0.3/). Although comprehensive testing coverage does not guarantee that the package is bug-free, it eliminates obvious errors, e.g. accessing local variables before definition.

Comprehensive unit testing greatly helps with validating correctness, ensuring reproducibility and refactoring the code. We carefully tested our analytical procedures with multiple input datasets to cover standard and edge cases, in order to make sure that the results are correct and reproducible. We also confidently refactored the code multiple times throughout the development process to improve backend performance, accommodate special use cases, and reorganize intra-package dependencies. For a non-trivial package with multiple interrelated components like scedar, changes in certain components may unexpectedly affect other components that directly or indirectly use the changed ones, so that validating the correctness after the changes would require a significant amount of effort without the comprehensive tests we deliberately built into scedar.

#### 1.2 Minimum description length

Minimum description length (MDL) is the minimum information size required to describe a set of data by a model (Hansen and Yu 2001). If the model plainly describes the data verbatim, the MDL of the data is equivalent to the size of the data. The MDL of the data can be reduced by a more sophisticated model that exploits the statistical properties in the data. For example, if 95% of the entries in a 10,000 x 10,000 matrix are 0s, a model can record only the indices of non-0 entries and keep a note that all other entries are 0s, which would be able to greatly reduce the MDL of the matrix. However, we do not want the model to become so complex that the information size for describing the model is very large. For example, a sequence of 10 single digit decimal integers can be described by a n=1-100 index model that stores all possible sequences, which is clearly larger than plainly describing the original 10 integers, and thus would not be a good model in the MDL framework. The principle of MDL is applied in statistics and machine learning to select the model that requires the smallest size of information to describe the model and data, and the practice is reviewed by Hansen and Yu in detail (Hansen and Yu 2001).

Using the principle of MDL, we developed a single-cell clustering method for scRNA-seq data, MIRAC. In the context of single-cell analysis, MIRAC finds the partition of cells (columns) in the matrix that yields the shortest code length of the data. The input data of MIRAC from scRNA-seq data could be any $n \times m$ matrix $X$ with $n$ cells (columns) and $m$ features (rows), where the features could be any measurements containing information about the similarity between the cells, e.g. the number of reads mapped to certain genes, distances to certain cells from a common coordinate, or dimensionality reduced coordinates in a feature space. In order to code the $n \times m$ data matrix $X$, we use a two-stage scheme (Hansen and Yu 2001), in which we code the partition of cells in the first stage and the partitioned data in the second, which we discuss in the following sections.

##### 1.2.1 Minimum description length in practice

Practically, let an observation $x$ of a random variable $X$ follow an arbitrary probability distribution $P$ with parameters $\Theta = \{\theta_1, \theta_2, ..., \theta_k\}$. If $P$ is continuous, let $f$ be its probability density function, otherwise probability mass function. Then, the code length of $x$ is $-\log f(x)$ with an arbitrary base of $2$ or the Euler's number $e$. In scedar, we consistently use $e$ as the base. The code length of $n$ observations $\{x_1, x_2, ..., x_n\}$ is the sum of the code lengths of individual observations, which gives an overall code length of $\sum_{i=1}^{n} -\log f(x_i)$. The code length of $P$ is the code length of $\Theta$ using uniform distribution for each parameter.

In a two-stage coding scheme, the overall code length of the data, i.e. the observations, is the sum of the following:

- Stage 1: code length of the statistical model $\mathcal{M} = \{P, \Theta\}$ (for the observations).
- Stage 2: code length of the observations encoded using $\mathcal{M}$.

When there are multiple statistical models $\{\mathcal{M}_1, \mathcal{M}_2, ..., \mathcal{M}_t\}$, we select the one that gives the shortest overall code length of the data. Intuitively, the closer the assumed distribution $P$ to the true distribution of the observations, the shorter the overall code length of the observations. The simpler the assumed distribution $P$, the shorter the code length of $P$.

Importantly, the code length of observations encoded by a discrete model cannot be directly compared to the code length of the same observations encoded by a continuous model. For example, let $X = [1, 1, 0, 1, 0]$ be our observations. The code length of $X$ encoded by $Bernoulli(0.6)$ is $L(X, Bernoulli(0.6)) = -3\log 0.6 - 2\log 0.4 \approx 3.365$. The code length of $X$ encoded by $Uniform(0, 1)$ is $L(X, Uniform(0, 1)) = -3\log 1 - 2\log 1 = 0$. Although $Bernoulli(0.6)$ better describes $X$ than $Uniform(0, 1)$, $L(X, Bernoulli(0.6)) > L(X, Uniform(0, 1))$.

The theoretical background of MDL is extensively reviewed by Hansen and Yu (Hansen and Yu 2001).

#### 1.2.2 Two-stage coding scheme for clustered scRNA-seq data

In order to clarify the coding scheme for scRNA-seq clustering analysis, we introduce the following definitions and notations:

- Denote an *ordered list* $Z$ of $n$ items as $\langle z_1, z_2, ..., z_n \rangle$.
- Define function $list(\{s_1, s_2, ..., s_n\}) = \langle s_1, s_2, ..., s_n \rangle$ to convert a set to a list.
- Let an $n \times m$ matrix $X$ be the data matrix of $n$ cells and $m$ features. We define the following operations:
    - $X_{i,\cdot}$ gives the $i$th row of $X$.
    - $X_{\cdot,j}$ gives the $j$th column of $X$.
    - gives the entry of $X$ at $i$th row and $j$th column.
    - $X_{\langle i_1, i_2, ..., i_r \rangle, \langle j_1, j_2, ..., j_c \rangle}$ gives a matrix of crossed entries of $\langle i_1, i_2, ..., i_r \rangle$ rows and $\langle j_1, j_2, ..., j_c \rangle$ columns in $X$.
- Define a *partition* $P$ of a set $S$ as a set of non-empty subsets of $S$ that are disjoint, of which the union is the same as $S$. For example, $\{\{1\}, \{2, 3\}\}$ is a partition of $\{1, 2, 3\}$, whereas $\{\{1\}, \{2\}\}$ or $\{\{\}, \{1, 2, 3\}\}$ is not a partition of $\{1, 2, 3\}$.
- Define operation $|S|$ on any set $S$ to give the number of elements in $S$, i.e. cardinality of $S$.
- We use a partition of a set of $n$ different integers to denote a possible clustering result of $n$ cells.
- For any partition $P = \{P_1, P_2, ..., P_k\}$ of $S$:
    - $P$ is a *singleton partition* if $k = 1$.
    - We call $P_i$ the $i$th *cluster*, where $i \in \{1, 2, ..., k\}$.
    - We define function $I(i, P)$ on any element $i \in S$, and $I(i, P) = j$ such that $i \in P_j$. Thus, we have a pair $(i, I(i, P))$ for each element $i \in S$, and we call $I(i, P)$ as the *cluster label* of $i$.
    - Let $B$ be the list of cluster labels $\langle I(1, P), I(2, P), ..., I(n, P) \rangle$.

For any $X_{n \times m}$, $P = \{P_1, P_2, ..., P_k\}$, and cluster labels $B$, we encode $X$ in the following two stages:

- Encode $B$ using categorical distribution. The code length of $B$ is

$$L(B) = \sum_{i=1}^{k} -|P_i| \log \frac{|P_i|}{n}.$$

- Encode $X$ as $\{X_{\text{list}(P_1),\cdot}, X_{\text{list}(P_2),\cdot}, ..., X_{\text{list}(P_k),\cdot}\}$. Within each row subset $X_{\text{list}(P_z),\cdot}$ of $X$, $m$ features are coded as individual random variables following arbitrary distributions. The code length of $X_{\text{list}(P_z),\cdot}$ is

$$L(X_{\text{list}(P_z),\cdot}) = \sum_{j=1}^{m} \sum_{i \in P_z} -\log f_j(X_{i,j}),$$

where $f_j$ is the probability density or mass function of the assumed distribution of $X_{\text{list}(P_z),j}$.

We write the code length of $X$ with partition $P$ as $L(X, P)$. When $P$ is a singleton partition, we omit $P$ and write $L(X)$.

#### 1.3 Mathematical theories on high-dimensional data analysis

The following two mathematical results on high-dimensional data analysis guided our development of analytical methods for scRNA-seq datasets.

##### 1.3.1 Distances between points in high-dimensional space

As the number of features increases, all samples become closer in similarity metrics to each other (Domingos 2012), in a sense that that the distance between a sample and its nearest sample approaches to the distance between the sample and its farthest sample (Beyer et al. 1999; Aggarwal, Hinneburg, and Keim 2001). This property of distance in high-dimensional space is also called distance concentration effect (Zimek, Schubert, and Kriegel 2012). Therefore, analytical methods based on distances, such as hierarchical agglomerative clustering, are less stable or, in other words, more susceptible to noise in the data. This result is mathematically described in the context of the nearest neighbors of a query point as following.

Definitions:

- Let any positive integer $m$ be the variable that the distance distributions may converge under. The variable $m$ can be interpreted as dimensionality, but this interpretation is not required by the proof of Theorem 1 given by Beyer *et al.* (Beyer et al. 1999).
- Let $n$ be the number of points.
- Let $X_1^m, X_2^m, ..., X_n^m$ be $n$ independent points such that $X_i^m \sim P_X^m$ for any $i \in \{1, 2, ..., n\}$, where $P_X^m$ a probability distribution.
- Let $Q^m$ be a query point sampled from the probability distribution $P_Q^m$ independently from $X_1^m, X_2^m, ..., X_n^m$.
- Let $0 < p < \infty$ be a constant.
- Define $D_m(X_i^m, Q^m)$ for any $i \in \{1, 2, ..., n\}$ as a function that returns a non-negative real number.

- Denote $D_m^{\min} = \min\{D_m(X_1^m, Q^m) | i \in \{1, 2, ..., n\}\}$.
- Denote $D_m^{\max} = \max\{D_m(X_1^m, Q^m) | i \in \{1, 2, ..., n\}\}$.

**Theorem 1.** *(Beyer et al.) If*

$$\lim_{m \to \infty} \mathrm{var}\left(\frac{(D_m(X_1^m, Q^m))^p}{E[(D_m(X_1^m, Q^m))^p]}\right) = 0 \ ,$$

*then for every* $\epsilon > 0$

$$\lim_{m \to \infty} P[D_m^{\max} \leq (1 + \epsilon) D_m^{\min}] = 1 \ .$$

The proof of Theorem 1 is given by Beyer *et al.* (Beyer et al. 1999). From the theorem, given that the distance distribution follows certain condition as $m$ increases, the distances of all points to the query point converges to a constant, which implies that the concept of nearest neighbor may not be meaningful (Beyer et al. 1999; Aggarwal, Hinneburg, and Keim 2001). The extent of restrictiveness of the precondition is also discussed by Beyer *et al*. (Beyer et al. 1999).

Although this property of distance between high-dimensional points affects analytical methods relying on distances (Beyer et al. 1999; Aggarwal, Hinneburg, and Keim 2001), the influences could be alleviated by dimensionality reduction, of which the performance in preserving the pairwise distances is characterized by the Johnson–Lindenstrauss lemma (Theorem 2) (Johnson and Lindenstrauss 1984).

#### 1.3.2 Johnson–Lindenstrauss lemma

The Johnson–Lindenstrauss lemma generally states that $n$ high-dimensional points in Euclidean space can be embedded into a lower dimensional Euclidean space with $O(\log n / \epsilon^2)$ dimensions for any $0 < \epsilon < 1$, while preserving the pairwise distances between $n$ points with errors within a factor of $\epsilon$ (Dasgupta and Gupta 1999; Johnson and Lindenstrauss 1984). The mathematical description of the theorem is summarized by Dasgupta and Gupta (Dasgupta and Gupta 1999) as the following:

***Theorem 2.*** *(Johnson–Lindenstrauss lemma) For any* $0 < \epsilon < 1$ *and any integer* $n$*, let* $k$ *be a positive integer such that*

$$k \geq 4(\epsilon^2/2 - \epsilon^3/3)^{-1} \ln n \ .$$

*Then for any set* $V$ *of* $n$ *points in* $\mathbb{R}^d$*, there is a map* $f : \mathbb{R}^d \to \mathbb{R}^k$ *such that for all* $u, v \in V$*,*

$$(1 - \epsilon) u - v^2 \leq f(u) - f(v)^2 \leq (1 + \epsilon) u - v^2 \ .$$

*Further this map can be found in randomized polynomial time.*

The proof of the Johnson–Lindenstrauss lemma with elementary probabilistic techniques is given by Dasgupta and Gupta (Dasgupta and Gupta 1999).

#### 1.4 Skewed root division of a hierarchical agglomerative clustering tree

We implemented a simple procedure to skew a hierarchical agglomerative clustering (HAC) before dividing the root into left and right subtrees (**Figure S10**). The skewed tree ensures that the smaller subtree of the root has $\geq n_{min}^{cluster}$ leaves, while preserves the ordering of leaves and maintains the invariants of a HAC tree. When $n_{min}^{cluster}$ is equal to half of the number of all leaves, the resulting division is similar to a balanced one.

Skewed root division is optional in Minimum description length (MDL) iteratively regulated agglomerative clustering (MIRAC). The procedure could be used to ensure that the divided sub-clusters are not too small, in order to improve the robustness of MDL estimation, because MDL estimation of a sub-cluster of too few samples is susceptible to noise.

**input :** $X$, a numeric transcription level matrix of size $n \times m$

$S$, a numeric pairwise similarity matrix of size $n \times n$

$x^{\mathsf{min}}$, minimum of an entry in $X$ to be considered as transcribed

$k$, the number of nearest neighbors to check for picking up gene dropouts

$n^{\mathsf{dropout}}$ ($\leq k$), for a zero entry $X[i, j]$ to be called a dropout, the minimum number of cells in $k$ nearest neighbors of cell $i$ transcribing the gene $j$

$n^{\mathsf{iter}}$, the number of iterations

**output:** $A$, a numeric matrix of size $n \times m$ storing the transcription levels of genes picked up

$A \leftarrow$ `zeros` $(n, m)$ $E \leftarrow X \geq x^{\mathsf{min}}$ `//` $E[i, j]$ `stores whether gene` $j$ `is transcribed in cell` $i$

**for** $i \leftarrow 1$ **to** $n^{\mathsf{iter}}$ **by** 1 **do**

  $n_i^{\mathsf{dropout}} \leftarrow n^{\mathsf{dropout}} + \lceil \frac{n^{\mathrm{iter}} - i}{n^{\mathrm{iter}}} (k - n^{\mathsf{dropout}}) \rceil$ `// dropout threshold at` $i$`th iteration`

  **for** $j \leftarrow 1$ **to** $n$ **by** 1 **do**

    $I_{j\mathsf{knn}} \leftarrow$ `kNNIndices` $(S, j)$

    **for** $g \leftarrow 1$ **to** $m$ **by** 1 **do**

      **if** $X[j, g] = 0$ **then** `// potential dropout`

        $n_{j\mathsf{knn}}^{\mathsf{exp}} \leftarrow$ `sum` $(E[\, I_{j\mathsf{knn}} \,, g])$

        **if** $n_{j\mathsf{knn}}^{\mathsf{exp}} \geq n_i^{\mathsf{dropout}}$ **then** `// pick up gene dropout`

          $A[j, g] \leftarrow$ `median` $(X[\, I_{j\mathsf{knn}} \,, g])$

          $X[j, g] \leftarrow A[j, g]$

          $E[j, g] \leftarrow A[j, g] \geq x^{\mathsf{min}}$

        **end**

      **end**

    **end**

  **end**

**end**

**Algorithm S1**. Impute gene dropouts by k-nearest neighbors

**input :** $S$, a numeric pairwise similarity matrix of size $n \times n$

$s^{\mathsf{sim}}$ (scalar), the minimum similarity of between two cells called similar

$k$, the minimum number of nearest neighbors that are similar to a non-rare cell

$n^{\mathsf{iter}}$, the number of iterations

**output:** $A$, an indicator vector of lefgth $n$ storing whether a cell is rare or not

$A \leftarrow$ `zeros` $(n)$

$s^{\mathsf{min}} \leftarrow$ `min` $(S)$ `//` $s^{\mathsf{min}}$ `is a scalar of the overall minimum of` $S$

$s^{\mathsf{max}} \leftarrow$ `max` $(S)$ `//` $s^{\mathsf{max}}$ `is a scalar of the overall maximum of` $S$

**if** $s^{\mathsf{sim}} \leq s^{\mathsf{min}}$ **then return** $A$ `// all cells are similar`

**else if** $s^{\mathsf{sim}} > s^{\mathsf{min}}$ **then return** $A + 1$ `// all cells are not similar`

**for** $i \leftarrow 1$ **to** $n^{\mathsf{iter}}$ **by** $1$ **do**

  $s_i^{\mathsf{sim}} \leftarrow s^{\mathsf{sim}}$- $\frac{n^{\mathrm{iter}}-i}{n^{\mathrm{iter}}}(\ s^{\mathsf{sim}}\ -\ s^{\mathsf{min}}\ )$ `// minimum similarity threshold at` $i$`th iteration`

  **for** $j \leftarrow 1$ **to** $n$ **by** $1$ **do**

    **if** $A[j] \neq 1$ **then** `// cell` $j$ `is not marked as rare yet`

      $s_{j,k} \leftarrow$ `kthNNSimilarity` $(S,\ j,\ k,\ A)$`// ignore rare cells`

      **if** $s_{j,k} < s_i^{\mathsf{sim}}$ **then** $A[j] = 1$ `// mark cell` $j$ `as rare`

    **end**

  **end**

**end**

**return** A

**Algorithm S2**. Detect rare transcriptomic profiles by k-nearest neighbors

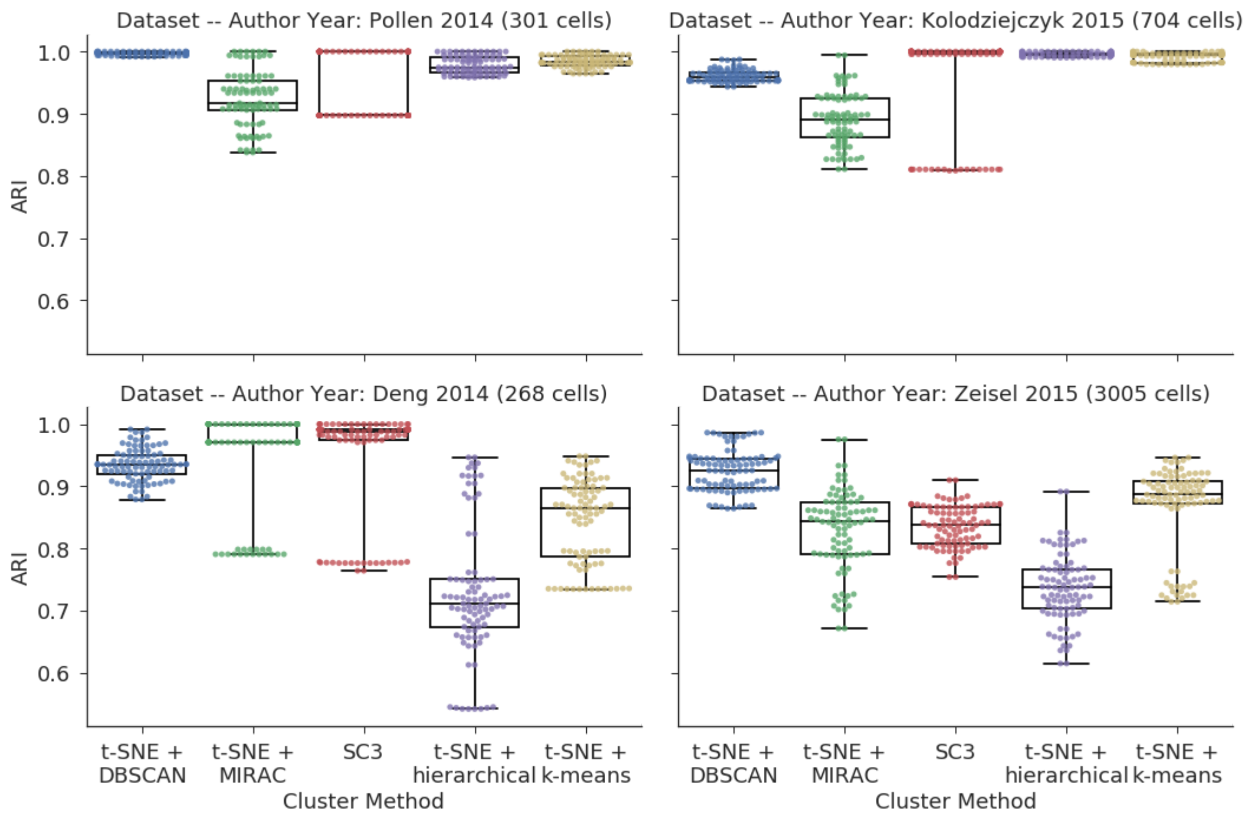


**Figure S1.** Stability of clustering methods on experimental dataset. Similarity between clustering results generated with different random states but the same parameters, quantified by adjusted rand index (ARI) (Hubert and Arabie 1985).

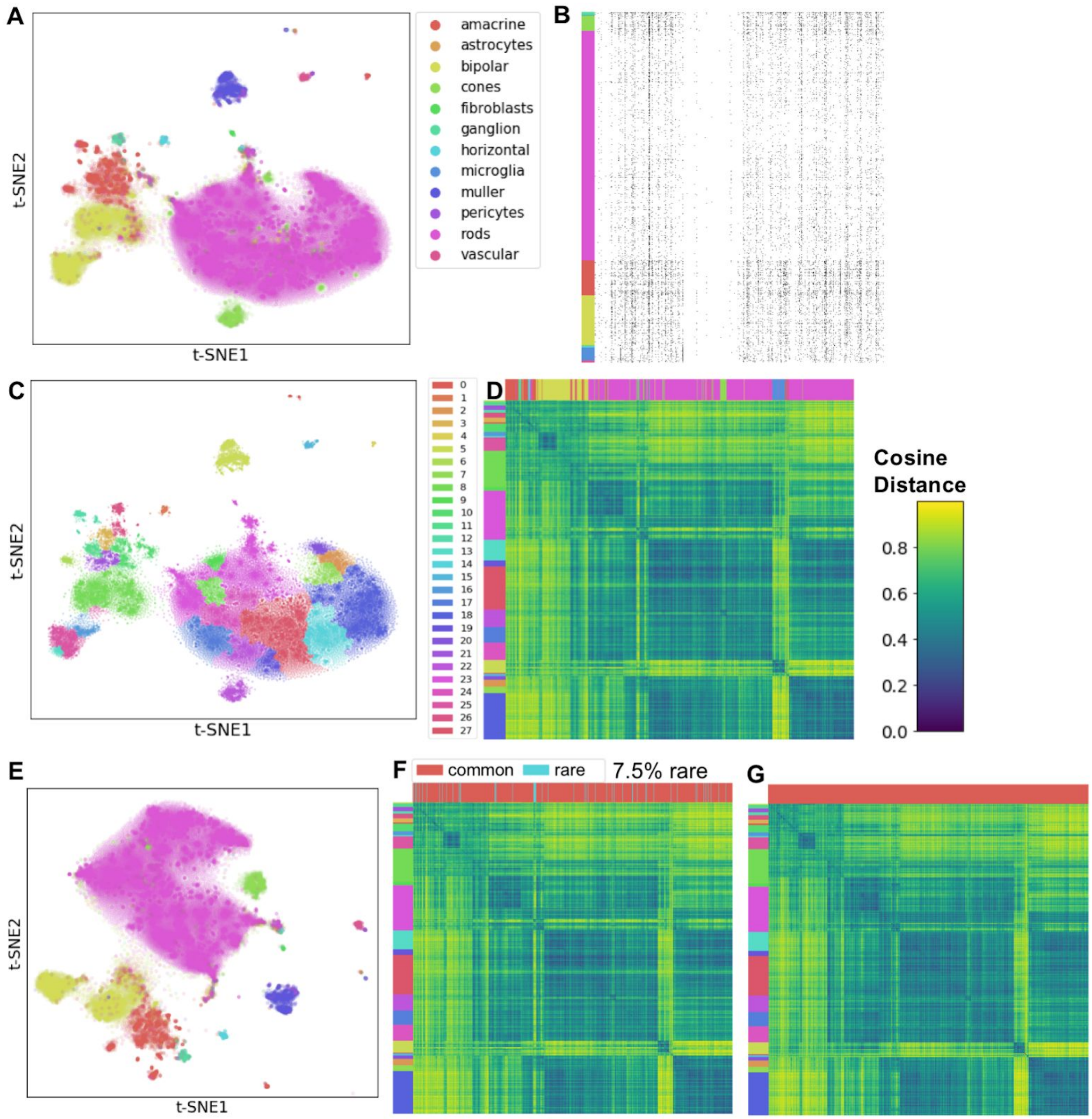


**Figure S2**. Scedar analysis of the scRNA-seq dataset containing 44,808 mouse retina cells generated by Drop-seq platform published by Macosko *et al*. (Macosko et al. 2015). (**A**) t-SNE scatter plot with cell type labels. (**B**) Read count matrix heatmap with rows as cells, columns as genes, and black color as ≥ 1reads. (**C**) t-SNE scatter plot with MIRAC labels. (**D**) Pairwise cosine distance heatmap with left strip as MIRAC labels and upper strip as cell type labels. (**E**) t-SNE scatter plot after KNN gene dropout imputation with cell type labels. (**F**) pairwise cosine distance heatmap with left strip as MIRAC labels and upper strip as common or rare transcriptomic profile labels. (**G**) pairwise cosine distance heatmap with rare transcriptomic profiles removed.

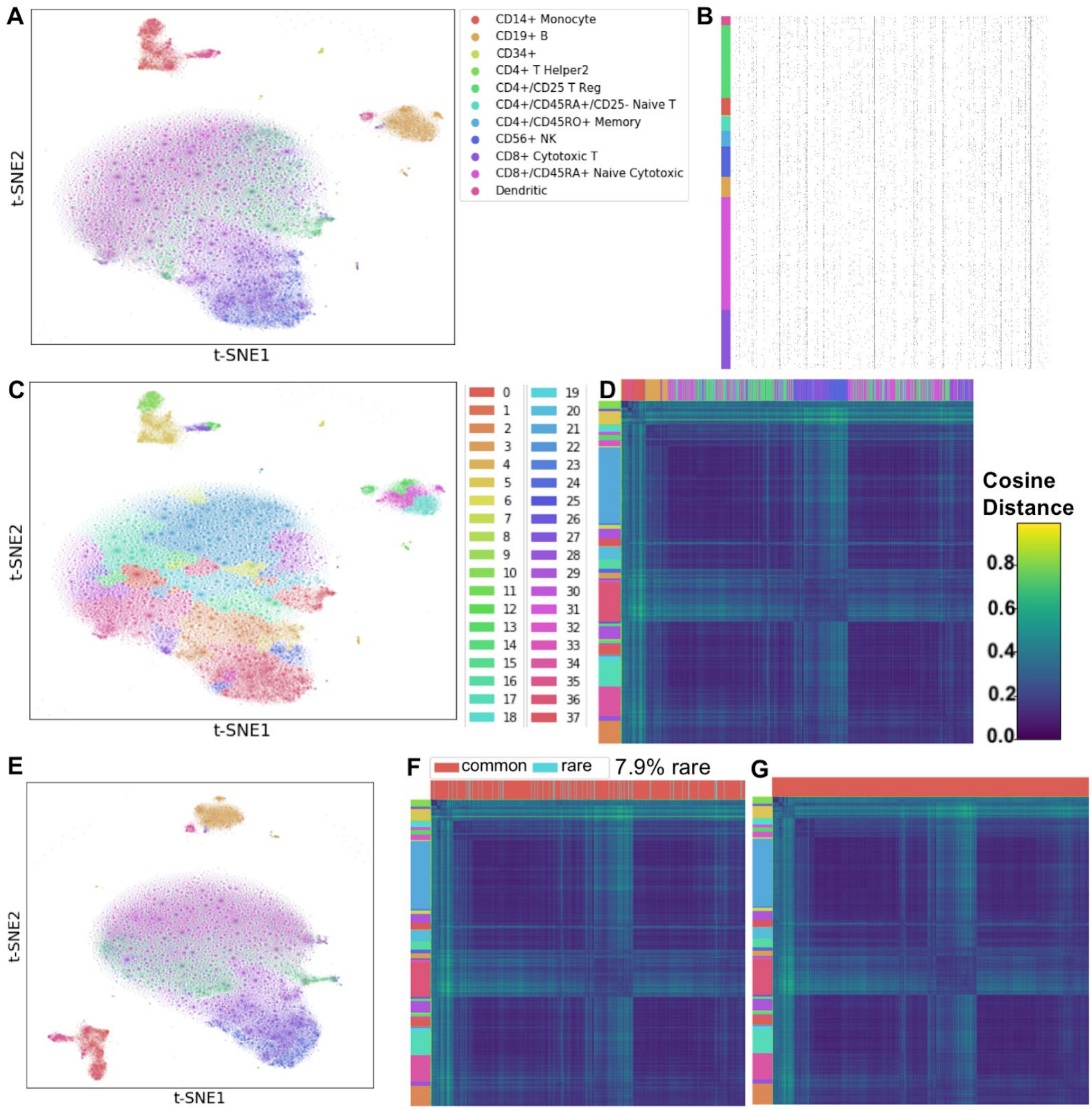


**Figure S3**. Scedar analysis of the scRNA-seq dataset containing 68,579 human peripheral blood mononuclear cells generated by 10x genomics GemCode platform published by Zheng *et al*. (Zheng et al. 2017). (**A**) t-SNE scatter plot with cell type labels. (**B**) Read count matrix heatmap with rows as cells, columns as genes, and black color as ≥ 1 reads. (**C**) t-SNE scatter plot with MIRAC labels. (**D**) Pairwise cosine distance heatmap with left strip as MIRAC labels and upper strip as cell type labels. (**E**) t-SNE scatter plot after KNN gene dropout imputation with cell type labels. (**F**) Pairwise cosine distance heatmap with left strip as MIRAC labels and upper strip as common or rare transcriptomic profile labels. (**G**) Pairwise cosine distance heatmap with rare transcriptomic profiles removed.

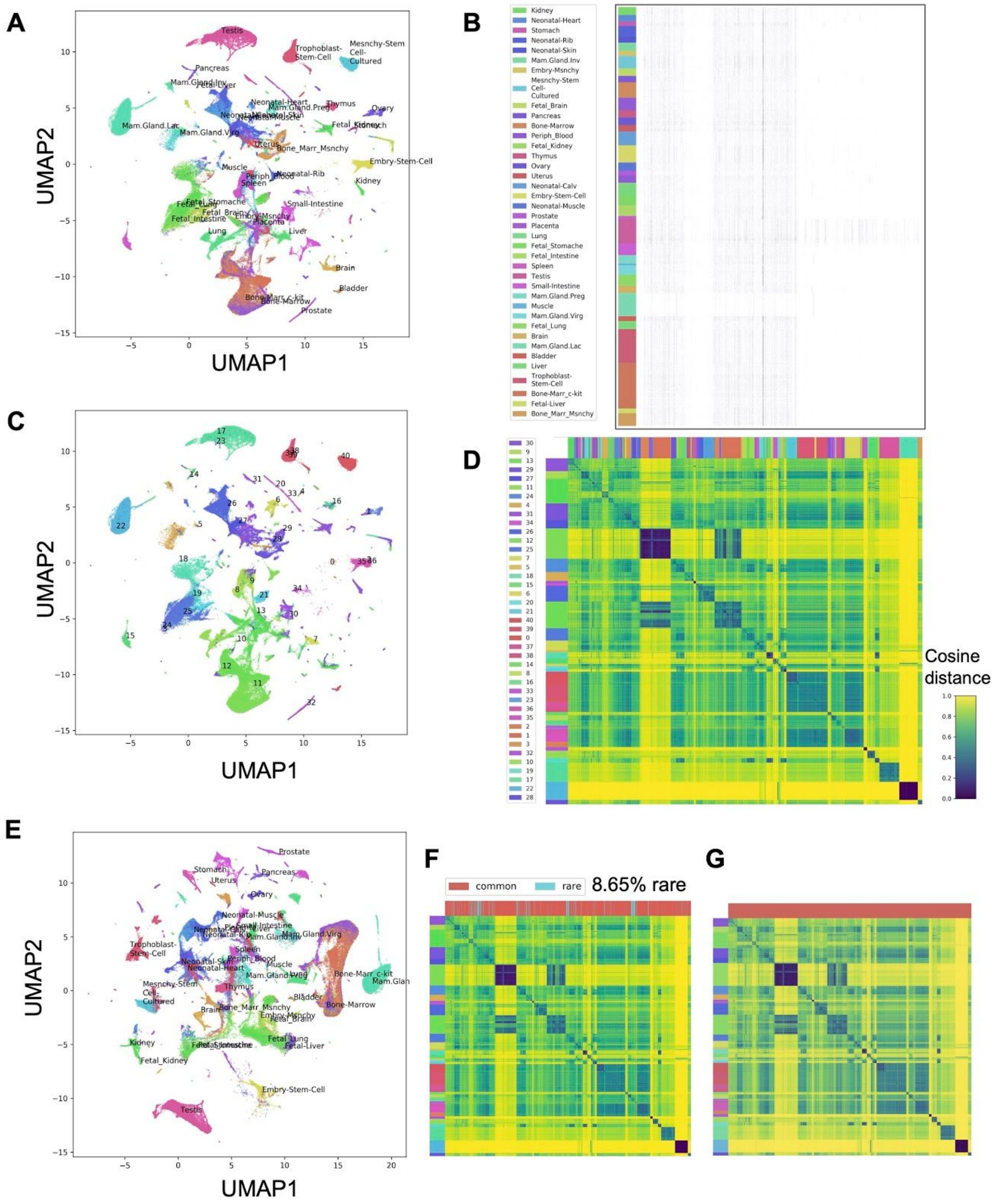


**Figure S4**. Scedar analysis of the scRNA-seq dataset containing 405,191 mouse cells from multiple tissues generated by Microwell-seq platform published by *Han et al.* (Han et al. 2018). (**A**) UMAP scatter plot with cell type labels. (**B**) Read count matrix heatmap with rows as cells, columns as genes, and black color as ≥ 1 reads. (**C**) UMAP scatter plot with MIRAC labels. (**D**) Pairwise cosine distance heatmap of downsampled 7449 cells with left strip as MIRAC labels and upper strip as cell type labels. (**E**) UMAP scatter plot after KNN gene dropout imputation with cell type labels. (**F**) Pairwise cosine distance heatmap of downsampled 7449 cells with left strip as MIRAC labels and upper strip as common or rare transcriptomic profile labels. (**G**) Pairwise cosine distance heatmap of downsampled 7449 cells with rare transcriptomic profiles removed.

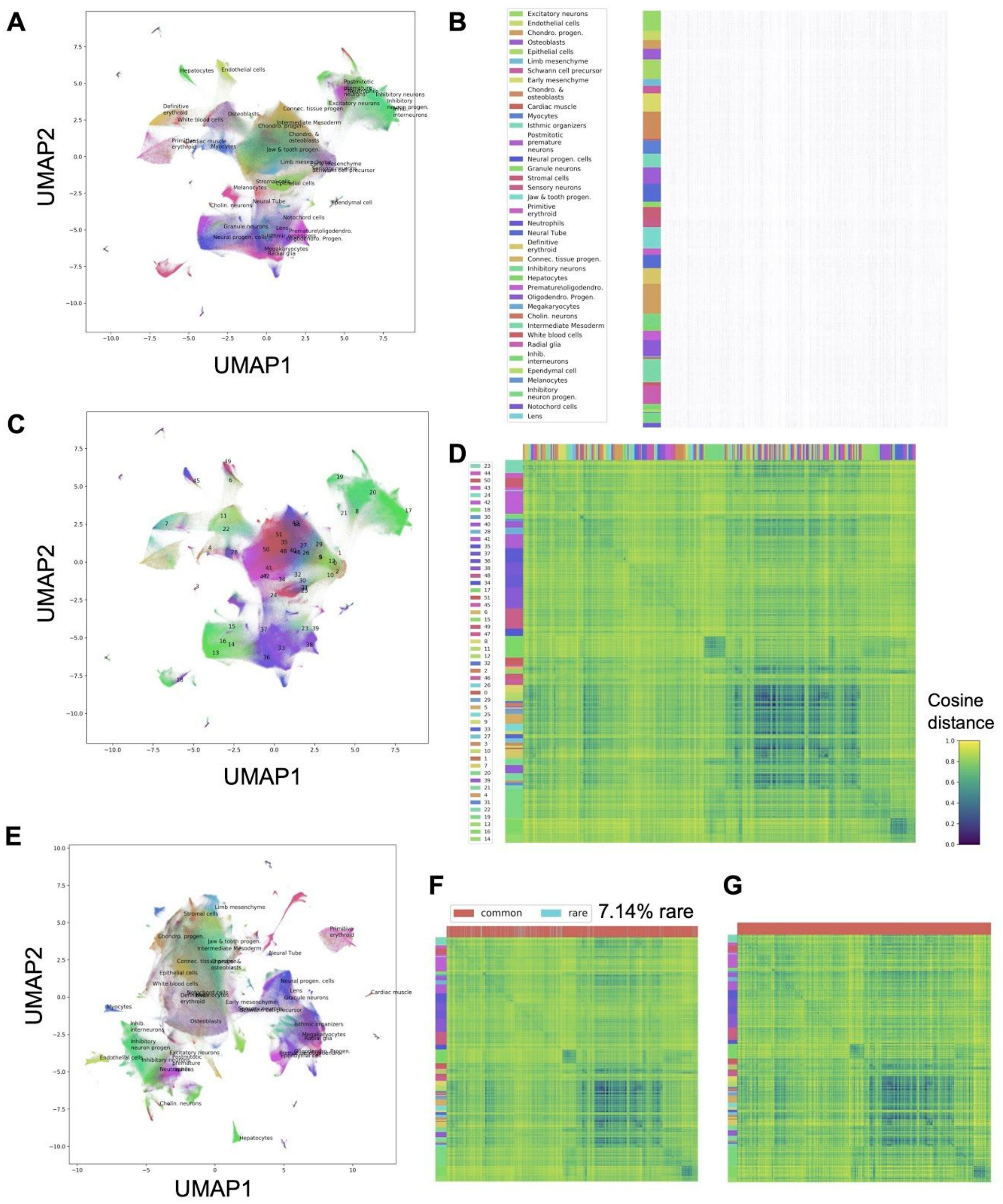


**Figure S5**. Scedar analysis of the scRNA-seq dataset containing 2,058,652 mouse cells from multiple tissues generated by sci-RNA-seq3 platform published by *Cao et al. (Cao et al. 2019)*. (**A**) UMAP scatter plot with cell type labels. (**B**) Read count matrix heatmap of downsampled 20,403 cells with rows as cells, columns as genes, and black color as ≥ 1 reads. (**C**) UMAP scatter plot with MIRAC labels. (**D**) Pairwise cosine distance heatmap of downsampled 20,118 cells with left strip as MIRAC labels and upper strip as cell type labels. (**E**) UMAP scatter plot after KNN gene dropout imputation with cell type labels. (**F**) Pairwise cosine distance heatmap of downsampled 20,118 cells with left strip as MIRAC labels and upper strip as common or rare transcriptomic profile labels. (**G**) Pairwise cosine distance heatmap of downsampled 20,118 cells with rare transcriptomic profiles removed.

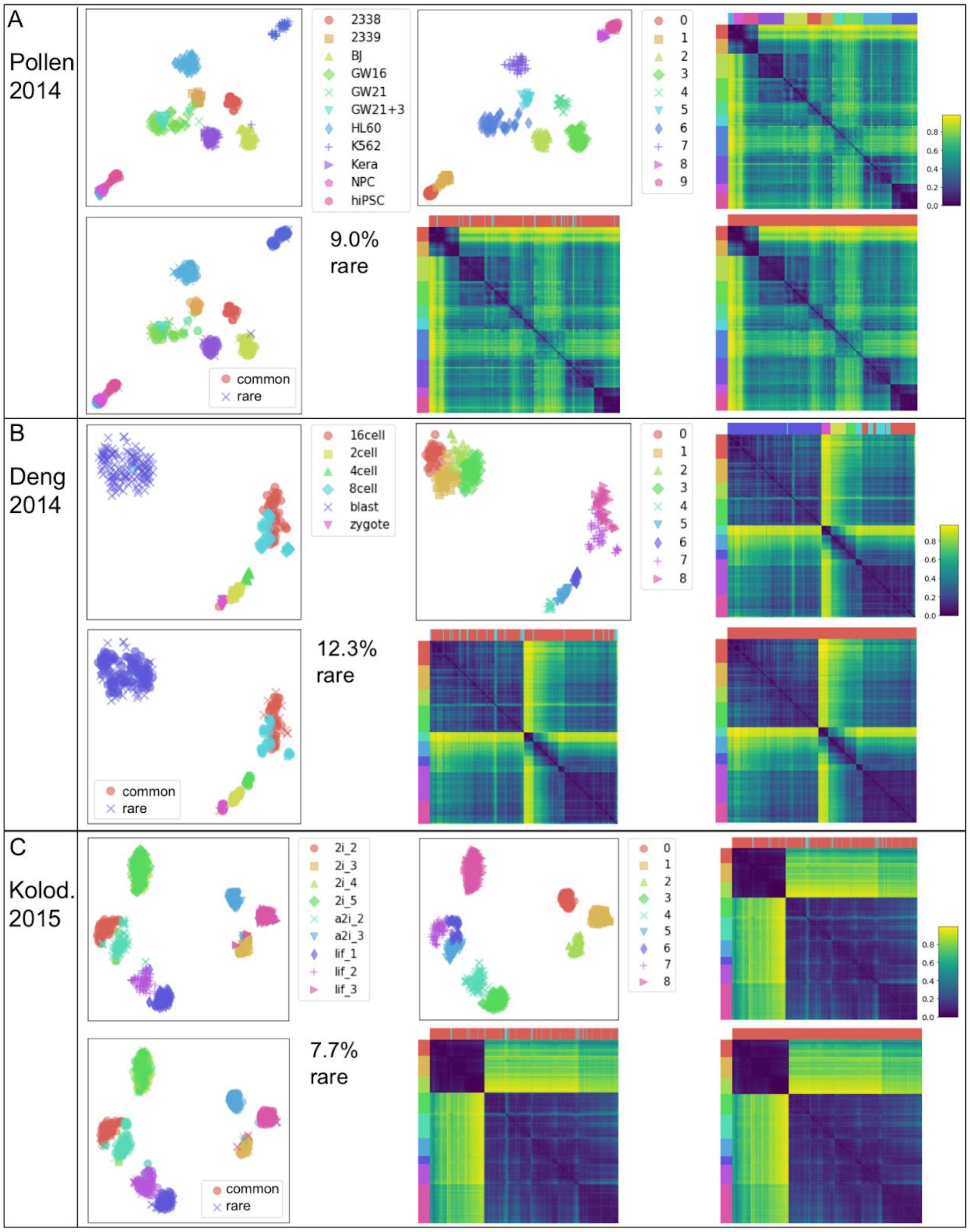


**Figure S6**. MIRAC and KNN rare transcriptomic profile detection results of experimental datasets. The sub-figures A, B, and C represent the results of scRNA-seq datasets published by Pollen *et al*. (Pollen et al. 2014), Deng *et al*. (Deng et al. 2014), and Kolodziejczyk *et al*. (Kolodziejczyk et al. 2015) respectively. Within each sub-figure, the plots are 1) t-SNE scatter plot with cell type labels, 2) t-SNE scatter plot with MIRAC cluster labels, 3) pairwise cosine distance heatmap with left strip as MIRAC labels and upper strip as cell type labels, 4) t-SNE scatter plot with common or rare rare transcriptomic profile labels, 5) pairwise cosine distance heatmap with left strip as MIRAC labels and upper strip as common or rare transcriptomic profile labels, 6) pairwise cosine distance heatmap with rare transcriptomic profiles removed.

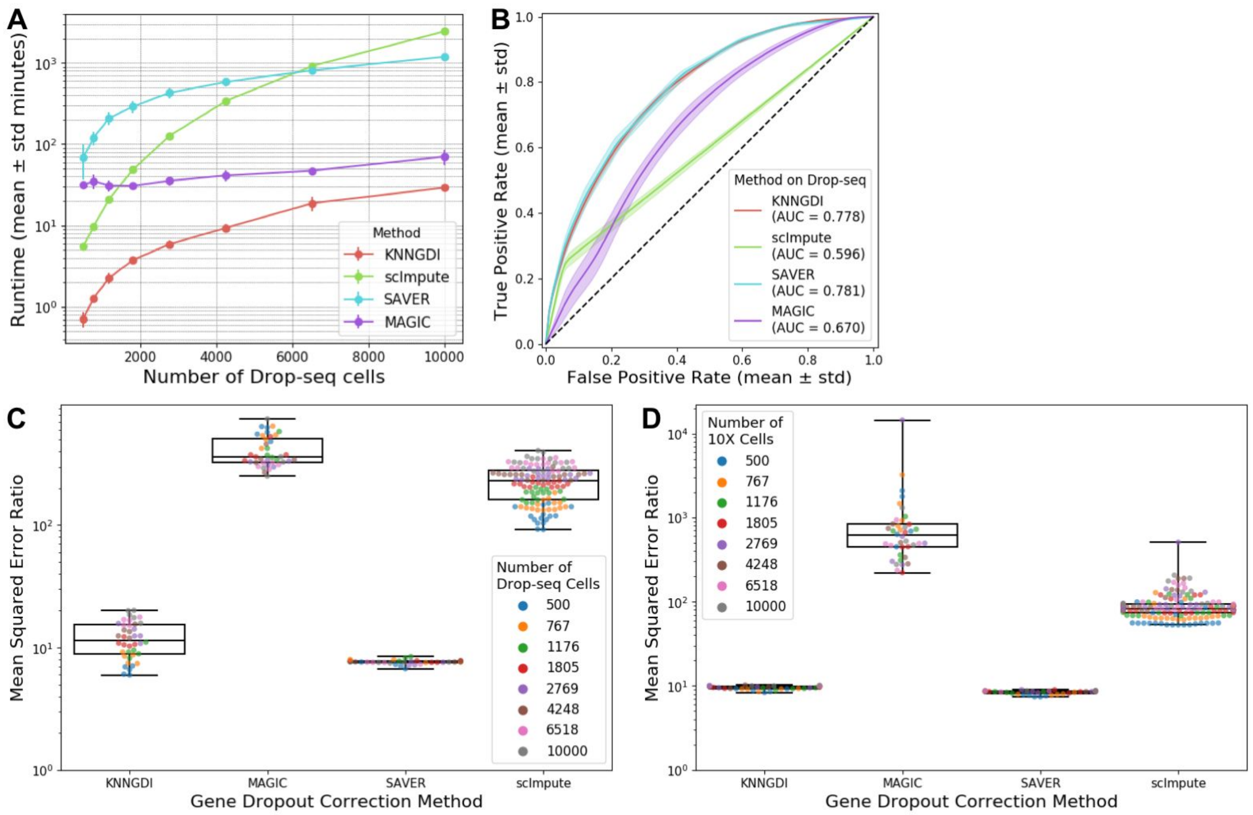


**Figure S7**. Benchmark results of gene dropout imputation methods. (**A**) Runtimes on 40 simulated Drop-seq datasets. (**B**) ROC curves (± standard deviation) of dropout detection on simulated Drop-seq datasets. (**C**) and (**D**) are mean squared error (MSE) ratios of different methods on simulated Drop-seq and 10x Genomics datasets respectively, where the MSE ratio is computed as the MSE of corrected read counts / MSE of true read counts.

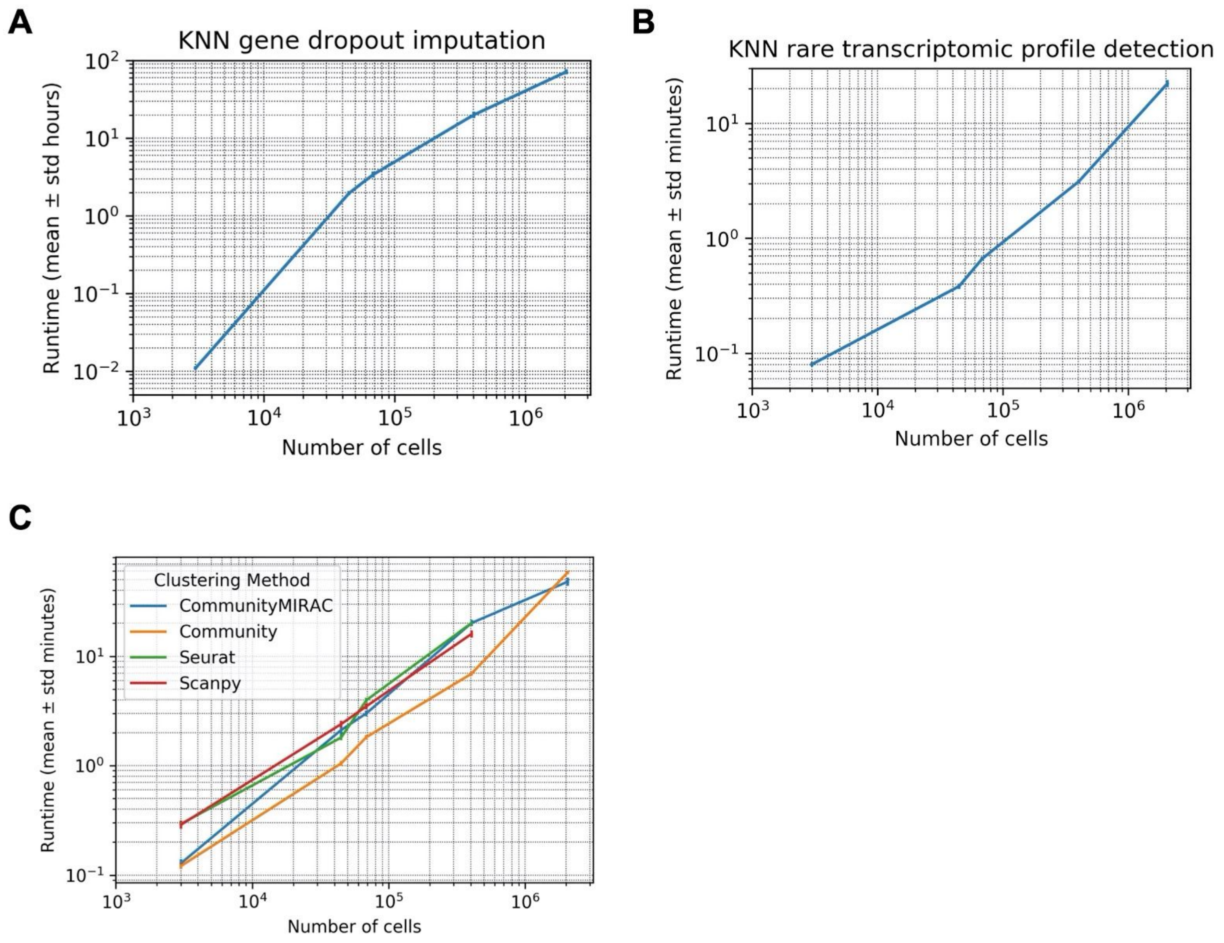


**Figure S8.** Runtimes of analytical methods implemented in scedar. **(A)** KNN gene dropout imputation. **(B)** KNN rare transcriptomic profile detection. **(C)** Community clustering, community extended MIRAC clustering, Seurat clustering, and Scanpy clustering. These methods were all performed on experimentally generated scRNA-seq datasets with 3005, 44808, 68579, 405191, and 2058652 cells (Zeisel et al. 2015; Macosko et al. 2015; Zheng et al. 2017; Han et al. 2018; Cao et al. 2019), except that Scanpy and Seurat were not able to cluster the mouse organogenesis cell atlas (MOCA) dataset that contain 2,058,652 single cells on a server with 1TB memory due to a mandatory conversion of the sparse read count matrix into dense matrix.

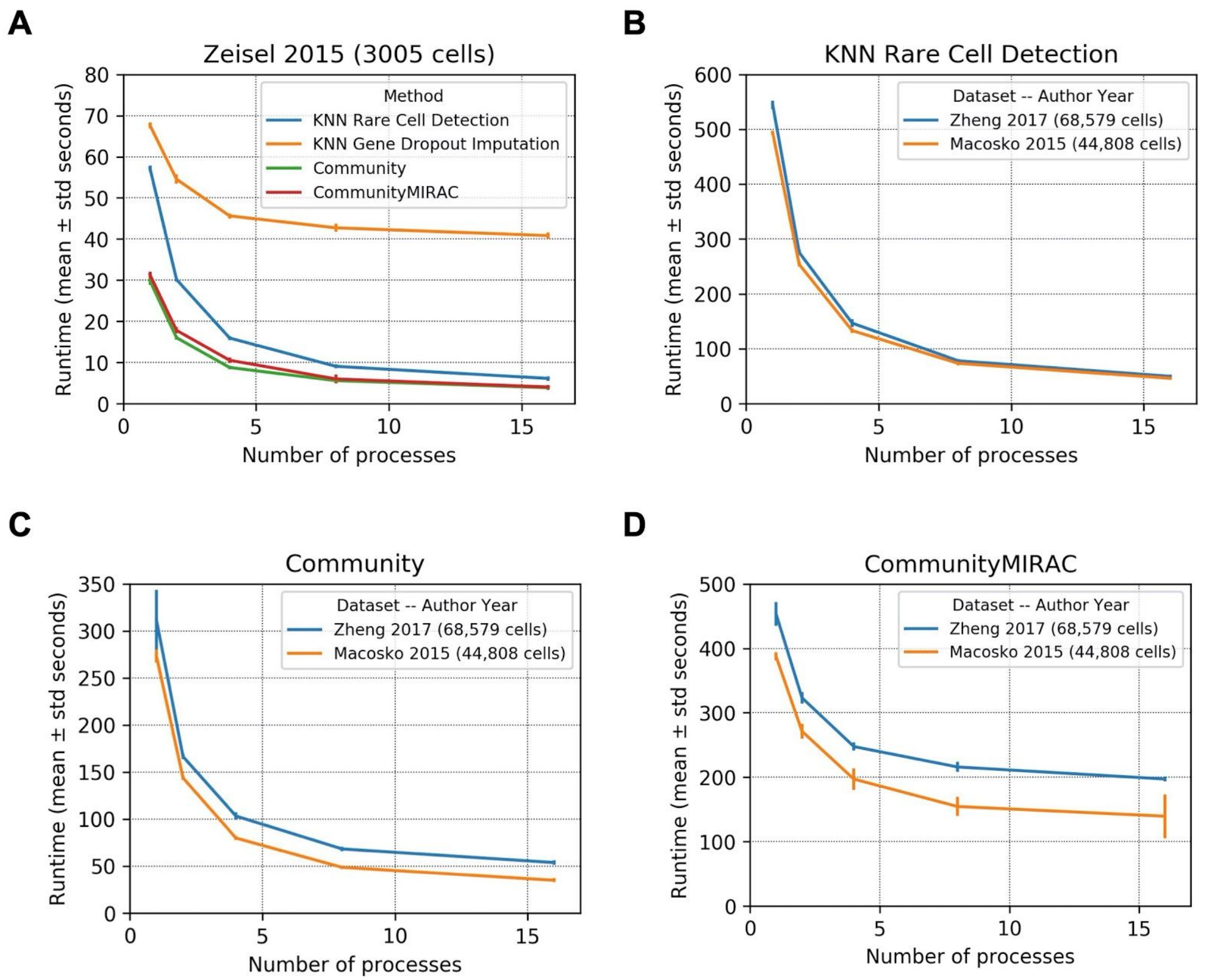


**Figure S9**. Runtimes of analytical methods implemented in scedar with different number of CPU cores for parallel computation. **(A)** All implemented methods performed on an scRNA-seq dataset with 3005 single cells (Zeisel et al. 2015). **(B)** KNN rare transcriptomic profile detection, **(C)** community clustering, and **(D)** community extended MIRAC clustering performed on two scRNA-seq datasets with 44,808 and 68,579 single cells (Zheng et al. 2017; Macosko et al. 2015).

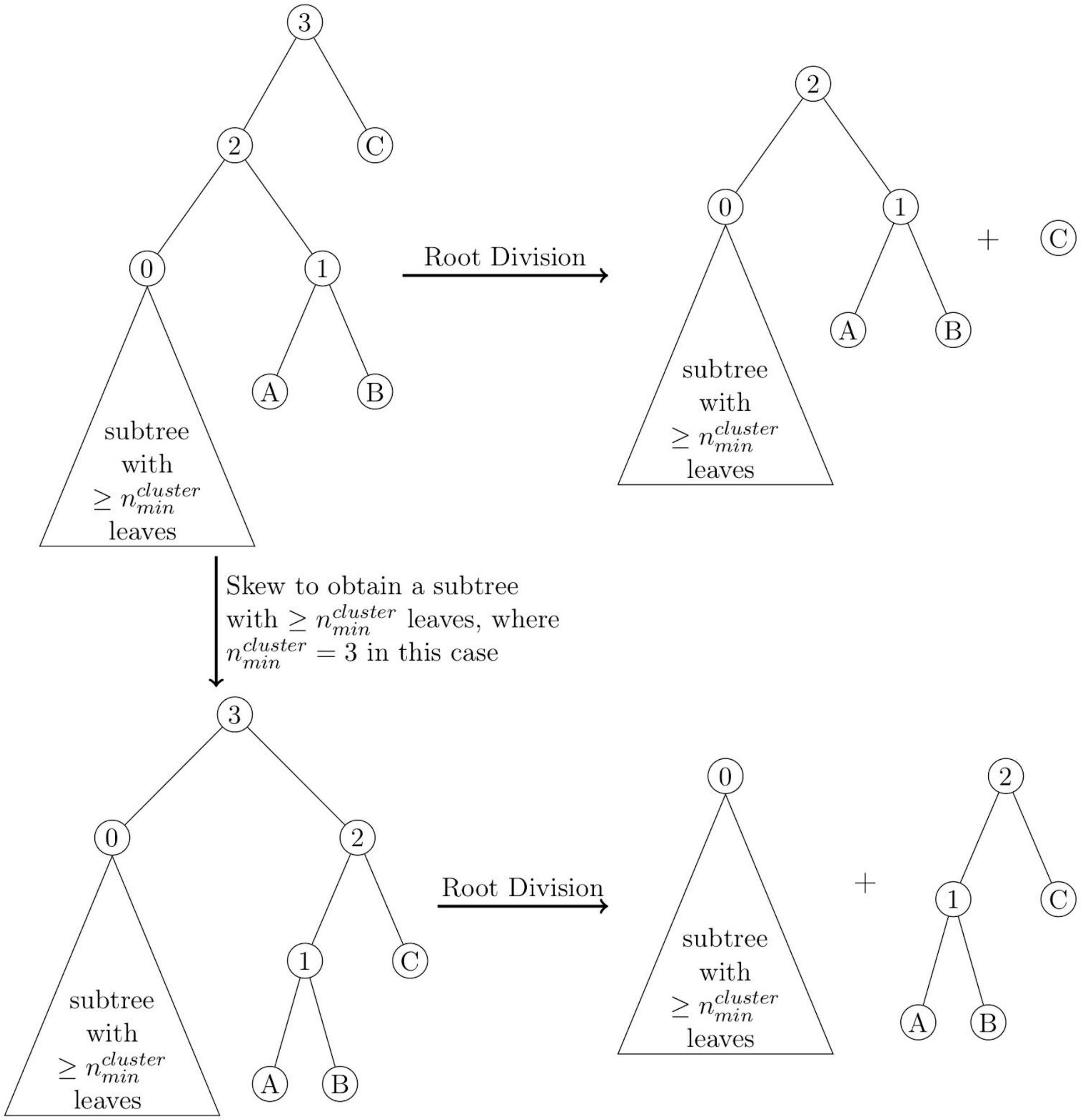


**Figure S10**. Skewed division of hierarchical agglomerative clustering tree. Tree leaves are samples, which are marked by upper case letters. Tree inner nodes are agglomerated samples by arbitrary linkage, which are marked by number. The triangle under inner node 0 represents an arbitrary valid subtree with $\geq n_{min}^{cluster}$ leaves. The root division procedure divides a tree into left and right subtrees of the root node. The skewing procedure creates a minimum subtree of the root with $\geq n_{min}^{cluster}$ leaves, where $n_{min}^{cluster} = 3$ in this specific case.

**Table S1.** Number of cell clusters for benchmark

| Publication | # cells | # published clusters | # cells per cluster | # benchmarked clusters |
|---|---|---|---|---|
| Deng *et al.* (2014) | 268 | 6 | 133, 50, 37, 22, 14, 12 | 6, 8, 9, 10, 11 |
| Pollen *et al.* (2014) | 301 | 11 | 54, 42, 40, 37, 26, 24, 22, 17, 17, 15 | 11, 12, 13, 14, 15 |
| Kolodziejczyk *et al.* (2015) | 704 | 9 | 93, 90, 82, 82, 81, 79, 72, 66, 59 | 9, 10, 11, 12, 14 |
| Zeisel *et al.* (2015) | 3005 | 9 | 948, 820, 390, 290, 198, 175, 98, 60, 26 | 15, 17, 20, 24, 30 |

**Table S2.** Top 20 Important Cluster ⟨1, 15, 22⟩ Separating Genes

| Gene | Importance | | # times used in 500 runs |
|---|---|---|---|
| | Mean | Std | |
| Mrc1 | 1.710 | 0.732 | 334 |
| F13a1 | 1.438 | 0.592 | 96 |
| Apoe | 1.420 | 0.579 | 438 |
| Pf4 | 1.385 | 0.625 | 91 |
| C1qb | 1.367 | 0.612 | 281 |
| Ms4a7 | 1.353 | 0.681 | 17 |
| Sepp1 | 1.333 | 0.471 | 3 |
| Dab2 | 1.333 | 0.471 | 3 |
| Cbr2 | 1.282 | 0.586 | 71 |
| Ctsc | 1.250 | 0.433 | 4 |
| Fcrls | 1.231 | 0.421 | 13 |
| Emr1 | 1.200 | 0.400 | 5 |
| Itm2a | 1.192 | 0.407 | 370 |
| Ccl24 | 1.158 | 0.446 | 152 |
| Fcgr3 | 1.143 | 0.389 | 140 |
| Cldn5 | 1.124 | 0.348 | 161 |
| C1qa | 1.071 | 0.258 | 14 |
| Sparc | 1.069 | 0.254 | 159 |
| Flt1 | 1.062 | 0.241 | 97 |
| Lyz2 | 1.059 | 0.235 | 17 |

**Table S3.** Software testing coverages of scRNA-seq data analysis tools. The tools are selected from the list curated by Zappia et al. (Zappia, Phipson, and Oshlack 2018).

| Package | Language | Package management | Test coverage | |
|---|---|---|---|---|
| | | | Qualitative | Quantitative |
| Scedar | Python | pip | comprehensive | 100% |
| slingshot | R | bioconductor | extensive | 87% |
| ZINB-WaVE | R | bioconductor | extensive | 81% |
| Scater | R | bioconductor | extensive | 69% |
| Splatter | R | bioconductor | extensive | 69% |
| BASiCS | R/C++ | bioconductor | extensive | 61% |
| TraCeR | Python | pip/conda | extensive | NA |
| Seurat | R/C++/Java | CRAN | extensive | NA |
| MAST | R | bioconductor | extensive | NA |
| Monocle | R/C++ | bioconductor | extensive | NA |
| PHATE | Python/R/Matlab | pip/cran/source | extensive | NA |
| Scanpy | python | pip | extensive | NA |
| pySCENIC | Python | pip | extensive | NA |
| scphaser | R | CRAN | extensive | NA |
| scran | R | source | extensive | NA |
| ZIFA | Python | source | extensive | NA |
| MAGIC | Python/R/Matlab | pip/cran/source | limited | NA |
| SCDE | R/c++/fortran | source | limited | NA |
| scLVM | Python/R | pip | limited | NA |
| scPipe | R | bioconductor | limited | NA |
| scruff | R | devtools source | limited | NA |
| SC3 | R/C++ | bioconductor | none | 0 |
| BackSPIN | Python | pip/conda | none | 0 |
| Mpath | R | source tarball | none | 0 |
| SAUCIE | Python | source | none | 0 |
| SAVER | R | bioconductor | none | 0 |
| scDD | R | bioconductor | none | 0 |
| GiniClust2 | R | source | none | 0 |
| SCnorm | R | bioconductor | none | 0 |

| | | | | |
|---|---|---|---|---|
| scMCA | R | source | none | 0 |
| scmap | R | source | none | 0 |
| SCODE | R | source | none | 0 |
| SCope | JavaScript/Python | npm | none | 0 |
| SCENIC | R | source | none | 0 |
| scImpute | R | source | none | 0 |
| scploid | R | source | none | 0 |
| Scrublet | Python | pip | none | 0 |
| scRutiNy | Python | pip | none | 0 |
| scTCRseq | Python | source | none | 0 |
| scTDA | Python | pip | none | 0 |
| SCUBA | Matlab | source | none | 0 |
| scVAE | Python | source | none | 0 |
| SIMLR | R | bioconductor | none | 0 |
| SINCERA | R | devtools source | none | 0 |
| SinQC | Python/R | source | none | 0 |
| SLICER | R | CRAN | none | 0 |
| SLICE | R | devtools source | none | 0 |
| SPADE | R | devtools source | none | 0 |
| StemID | R | devtools source | none | 0 |
| STEMNET | R | devtools source | none | 0 |
| URD | R | devtools source | none | 0 |
| Wishbone | Python | source | none | 0 |

McKinney, Wes, and Others. 2010. “Data Structures for Statistical Computing in Python.” In

*Proceedings of the 9th Python in Science Conference*, 445:51–56. Austin, TX.
Oliphant, Travis E. 2006. *A Guide to NumPy*. Vol. 1. Trelgol Publishing USA.
Pedregosa, Fabian, Gaël Varoquaux, Alexandre Gramfort, Vincent Michel, Bertrand Thirion, Olivier Grisel, Mathieu Blondel, et al. 2011. “Scikit-Learn: Machine Learning in Python.” *Journal of Machine Learning Research: JMLR* 12 (Oct): 2825–30.
Pollen, Alex A., Tomasz J. Nowakowski, Joe Shuga, Xiaohui Wang, Anne A. Leyrat, Jan H. Lui, Nianzhen Li, et al. 2014. “Low-Coverage Single-Cell mRNA Sequencing Reveals Cellular Heterogeneity and Activated Signaling Pathways in Developing Cerebral Cortex.” *Nature Biotechnology* 32 (10): 1053–58.
Traag, V. A., L. Waltman, and N. J. van Eck. 2019. “From Louvain to Leiden: Guaranteeing Well-Connected Communities.” *Scientific Reports* 9 (1): 5233.
Virtanen, Pauli, Ralf Gommers, Evgeni Burovski, Travis E. Oliphant, David Cournapeau, Warren Weckesser, alexbrc, et al. 2018. *Scipy/scipy: Scipy 1.1.0rc1*. https://doi.org/10.5281/zenodo.1218715.
Waskom, Michael, Olga Botvinnik, Drew O’Kane, Paul Hobson, Saulius Lukauskas, David C. Gemperline, Tom Augspurger, et al. 2017. *Mwaskom/seaborn: v0.8.1 (September 2017)*. https://doi.org/10.5281/zenodo.883859.
Zappia, Luke, Belinda Phipson, and Alicia Oshlack. 2018. “Exploring the Single-Cell RNA-Seq Analysis Landscape with the scRNA-Tools Database.” *PLoS Computational Biology* 14 (6): e1006245.
Zeisel, Amit, Ana B. Muñoz-Manchado, Simone Codeluppi, Peter Lönnerberg, Gioele La Manno, Anna Juréus, Sueli Marques, et al. 2015. “Brain Structure. Cell Types in the Mouse Cortex and Hippocampus Revealed by Single-Cell RNA-Seq.” *Science* 347 (6226): 1138–42.
Zheng, Grace X. Y., Jessica M. Terry, Phillip Belgrader, Paul Ryvkin, Zachary W. Bent, Ryan Wilson, Solongo B. Ziraldo, et al. 2017. “Massively Parallel Digital Transcriptional Profiling of Single Cells.” *Nature Communications* 8 (January): 14049.
Zimek, Arthur, Erich Schubert, and Hans-Peter Kriegel. 2012. “A Survey on Unsupervised Outlier Detection in High-Dimensional Numerical Data.” *Statistical Analysis and Data Mining* 5 (5): 363–87.
